# Sex-dependent macromolecule and nanoparticle delivery in experimental brain injury

**DOI:** 10.1101/817296

**Authors:** Vimala N. Bharadwaj, Connor Copeland, Ethan Mathew, Jason Newbern, Trent R. Anderson, Jonathan Lifshitz, Vikram D. Kodibagkar, Sarah E. Stabenfeldt

## Abstract

Development of effective therapeutics for brain disorders is challenging, in particular, the blood-brain barrier (BBB) severely limits access of the therapeutics into the brain parenchyma. Traumatic brain injury (TBI) may lead to transient BBB permeability that affords a unique opportunity for therapeutic delivery via intravenous administration ranging from macromolecules to nanoparticles (NP) for developing precision therapeutics. In this regard, we address critical gaps in understanding the range/size of therapeutics, delivery window(s), and moreover the potential impact of biological factors for optimal delivery parameters. Here we show, for the first time, to the best of our knowledge, that 24 h post-focal TBI female mice exhibit a heightened macromolecular tracer and NP accumulation compared to male mice, indicating sex-dependent differences in BBB permeability. Furthermore, we report for the first time the potential to deliver NP-based therapeutics within 3 d after focal injury in both female and male mice. The delineation of injury-induced BBB permeability with respect to sex and temporal profile is essential to more accurately tailor time-dependent precision and personalized nanotherapeutics.

## Introduction

Development of effective therapeutics for brain disorders, such as traumatic brain injury (TBI), remains a major clinical challenge^1–4^. In particular, the blood-brain barrier (BBB) severely limits access of the therapeutics into the brain parenchyma^1, 3^. Due to its highly selective nature, the BBB constitutes the greatest impediment for drug delivery via blood circulation to treat brain disorders. As such, several active drug delivery strategies such as transient BBB disruption via focused ultrasound and convection-enhanced delivery are being explored in preclinical settings to facilitate drug/macromolecules to cross or bypass the BBB^5^. These tools are useful, yet, certain neural pathologies may naturally afford the ability to exploit periods of enhanced BBB permeability for enabling drug delivery to the parenchyma. For example, TBI may lead to a transient BBB disruption, that affords a unique opportunity for the development of precision therapeutics via intravenous administration, thus facilitating a spectrum of therapeutics ranging from macromolecules to nanoparticles (NPs). However, there is a critical gap in understanding the permissible range/size of therapeutic agents, optimal delivery window(s), and moreover the potential impact of biological factors on BBB permeability after a TBI and ultimately therapeutic delivery parameters. Therefore, in this study, we characterize three crucial factors (1) size range (macromolecules vs NPs) for therapeutics, (2) their temporal profiles (acute and sub-acute post-injury delivery), and (3) sex dependence (female vs male) that can directly influence the accumulation within injured brain tissue.

Brain injury is characterized by structural failure and neurologic dysfunction that begins at the time of impact and lasting for hours to weeks^6, 7^. The primary injury is the direct result of the initial trauma of a TBI and leads to an indirect secondary injury, a cascade of neural and vascular events including inflammation, excitotoxicity, hypoxia, edema, and BBB disruption^6, 7^. The secondary injury develops over hours and days following a TBI allowing a time window for intervention. Most consequences of BBB disruption after brain injury are known to be detrimental, yet the disruption may also provide a transient window for delivery of therapeutics that normally would not cross this barrier from the systemic circulation^8–11^. Macromolecules such as specific antibodies and proteins have been identified as promising therapeutic agents for the treatment of various brain pathologies^1, 12^. These therapeutic agents can exert numerous biological actions in the brain such as regulation of cerebral blood flow, neurotransmission, and neuromodulation^12^. Furthermore, in recent years, NPs have been used extensively to serve as carriers for improved drug stability, pharmacokinetics, and therapeutic efficacy with reduced toxicity^13–19^. The above-mentioned classes of therapeutics including proteins^20, 21^ and NPs^22, 23^ have been approved by the US Food and Drug Administration for clinical use for other diseases/pathologies. However, there is a critical gap in understanding the utility of using both of these classes of therapeutics for brain injury since the delivery parameters for optimal and personalized delivery (i.e. the impact of biological factors) have not been previously investigated. Therefore, in this pre-clinical study, we use large molecular weight protein tracers (∼40 kDa, horseradish peroxidase) and NPs (∼40 nm, PEGylated fluorescent polystyrene nanoparticle) to address this critical gap and as a proof of concept to characterize the delivery profile to the injured brain tissue via the disrupted BBB.

Previously, we and others have established in both focal and diffuse injury models of TBI that macromolecular weight tracer (∼40 kDa)^19, 24–26^ and NPs^10, 19, 26–28^ robustly accumulate within injured brain tissue in the first 4h following TBI. Moreover, seminal TBI studies in male rodents suggest a biphasic BBB opening to macromolecular weight tracers with the first peak at acute (∼3-6h) time points followed by a delayed opening (∼3 d) after TBI^25, 29^. However, the potential to deliver larger therapeutics such as NPs beyond 24 h post-injury has not been investigated. Furthermore, sex differences are known to play a role in the morbidity and mortality after TBI^30, 31^, yet the underlying mechanisms are not well elucidated. Sex-specific preclinical studies have reported less BBB disruption in female rodents compared to males within 6h of injury^32, 33^ but no difference 24 h and 7 d post-injury^34, 35^. However, sex differences in BBB disruption during clinically important sub-acute time points between 24 h and 7 d after TBI have not been studied. Taken together, this study aims to characterize therapeutic window for delivery of large molecular weight tracer and NPs beyond 24 h post-injury. Moreover, we aim to evaluate potential sex-dependent differences in BBB disruption and subsequent large molecular weight tracer and NP accumulation following TBI. Ultimately, we reveal not only an extended NP delivery window but also key sex-dependent considerations for both large molecular weight tracer and NP delivery strategies following TBI.

## Results

The lateral controlled cortical impact (CCI) imparts an injury directly to the frontoparietal cortex generating a cortical lesion ipsilateral to the impact with no direct damage to the contralateral hemisphere. Brain injury may lead to a transient BBB disruption that could potentially allow therapeutic delivery to the injured brain regions. See Supplementary Fig S1 for experimental design/timeline. The focus of this study is three fold: (1) to investigate delivery dynamics of macromolecules and NPs, (2) to determine acute and sub-acute temporal delivery profiles, and (3) to examine sex dependence for delivery of these therapeutic agents.

### Extravasation of intravenously injected macromolecular tracer (horseradish peroxidase) after focal brain injury

To examine the potential delivery of the macromolecules, a large molecular weight (MW) tracer, horseradish peroxidase (HRP, 40 kDa), was used as a surrogate^36–38^. Each cohort received intravenous HRP injection 10 mins prior to sacrifice at 3 h, 24 h, 3 d, 7 d following CCI injury. The HRP extravasation was analyzed at the core of the injury (∼ −1.65 mm Bregma). The extravasation of HRP at 3 h, 24 h and 3 d time points was restricted to the primary injury region (Fig 1a). We observed no HRP staining in the contralateral hemisphere for any time point post-injury (Supplementary Fig S2). Two-way ANOVA analysis revealed a significant main effect of both hemisphere (p<0.0001) and time (p<0.0001) on HRP levels for both female and male cohorts (Fig 1b and 1c, respectively); a significant interaction between hemisphere and time was also discovered (p<0.0001) (Supplementary Table 1 for statistical details). Pair-wise analysis of extravasation of HRP in both female and male cohorts showed markedly higher levels of HRP stain at 3 h, 24 h, and 3 d post-injury compared to their respective contralateral hemispheres. At 7 d post-injury, the HRP staining in the ipsilateral location for both female and male cohorts was minimal and not significant compared to their contralateral hemisphere. At 3 h post-injury for both female and male cohorts, HRP staining was significantly different than the 24 h post-injury ipsilateral hemisphere. Interestingly, the male 3 d (ipsilateral) cohort showed a significant increase (∼150% increase) in HRP staining relative to the 24 h male ipsilateral group.

**Fig. 1:**
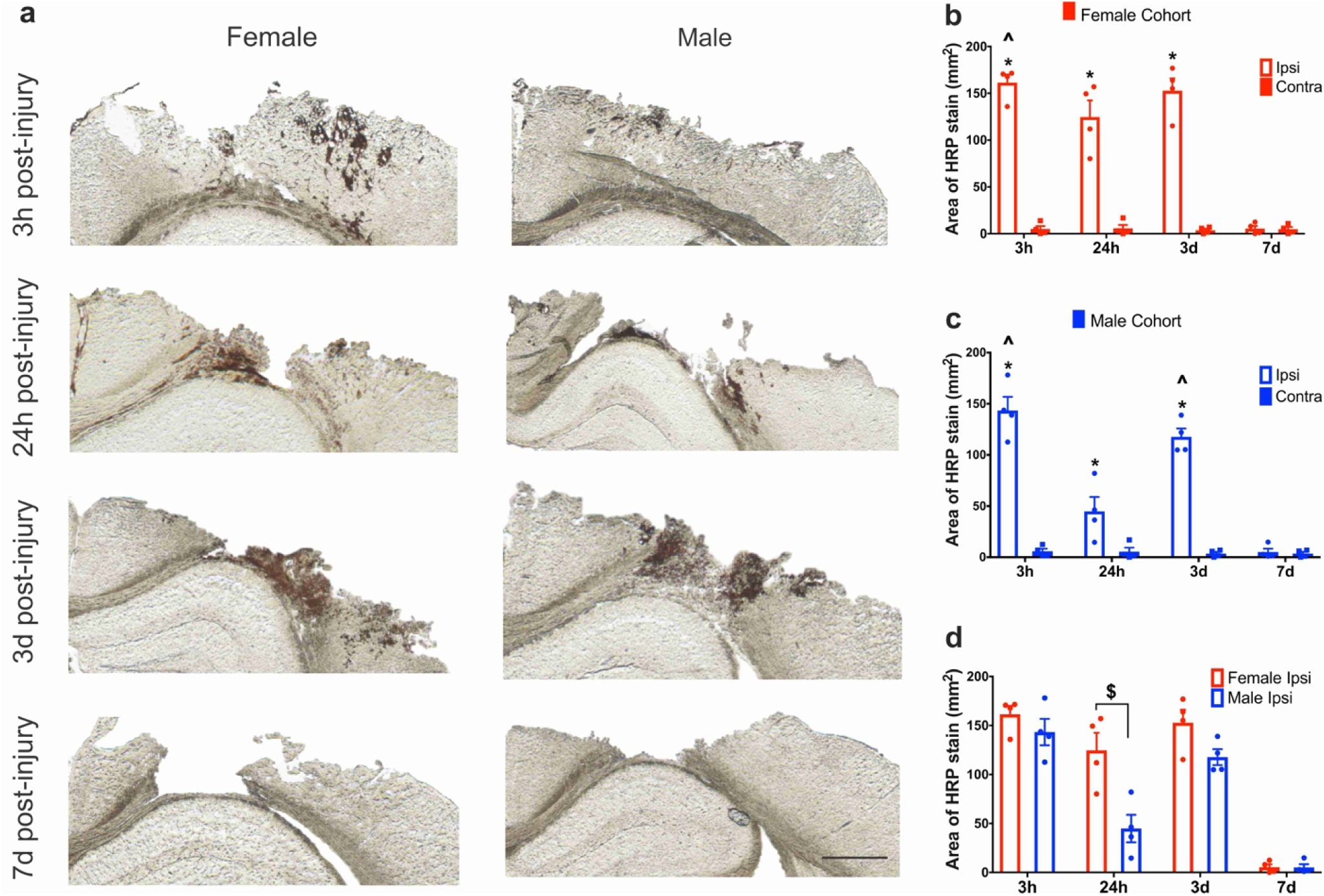
Macromolecular tracer (HRP) extravasation after CCI: **(a)** Ipsilateral hemisphere: Representative images of HRP extravasation (dark brown staining) at 3 h, 24 h, 3 d, and 7 d post-CCI. The first column shows the HRP response in the female cohort and the second column in males. Scale bar=1mm. (b-c) Sex dependence of HRP extravasation: Extravasation of HRP in the female cohort (a) and male cohort (b) at different time points post-injury. (d) Quantification of HRP stain in the ipsilateral hemisphere of female and male cohorts across different time points. *p<0.05 compared to their respective contralateral hemisphere and 7 d ipsilateral hemisphere, two-way ANOVA, Bonferroni’s multiple comparisons. ^ p<0.05 compared to 24 h ipsilateral hemisphere, $ p<0.05 compared to male 24 h ipsilateral hemisphere, two-way ANOVA, Bonferroni’s multiple comparisons. Error bars represent standard error of mean, n=4.

A two-way ANOVA revealed a significant main effect for both sex (p=0.0004) and time (p<0.0001) on ipsilateral HRP levels (Fig 1d); the interaction between sex and time was also significant (p=0.0119) (Supplementary Table 1 for statistical details). Specifically, at 24 h post-injury, there was significantly increased HRP staining in females compared (3 x higher) to their male counterparts (p= 0.0002). This observation supported the previously reported biphasic BBB disruption pattern for male cohorts^25, 29^. However, more intriguingly, the females exhibited consistent HRP staining levels at both 24 h and 3 d post-injury, not following a biphasic response. Taken together, the extravasation of the macromolecular tracer was observed acutely (3 h and 24 h) and sub-acutely (3 d) post-injury, demonstrating a potential for delayed therapeutic macromolecule delivery. Furthermore, more strikingly, the macromolecular tracer delivery was sex-dependent, where only the males showed a biphasic pattern.

### Accumulation of intravenously injected nanoparticles after focal brain injury

To examine the potential NP delivery at acute and sub-acute time points and the sex dependence for delivery after brain injury, we used fluorescent PEGylated 40 nm polystyrene NPs (See Supplementary Fig S3 for NP characterization). Each cohort received an intravenous injection of the NPs three hours before sacrifice (See Supplementary Fig S4 for biodistribution data). Mirroring HRP analysis, the area of NP accumulation was quantified at the core of the injury (∼ −1.65mm Bregma) via an epi-fluorescent microscope. NP accumulation was restricted to the primary injury region (Fig. 2 (a)), similar to the HRP staining. We observed no detectable NP accumulation in the contralateral hemisphere for any time point post-injury (See Supplementary Fig S5). Briefly, we found significant NP delivery for females acutely at 3 h and 24 h and sub-acutely at 3 d post-injury. However, for males, the NP accumulation was significantly increased in the ipsilateral hemisphere compared to the contralateral hemisphere only at 3 h for acute delivery and at 3 d for sub-acute delivery. Moreover, there was a significantly higher NP accumulation at 24 h post-injury in females compared to males (Fig. 2 (b); 2.5 x higher). These results trended similar to HRP extravasation.

**Fig. 2:**
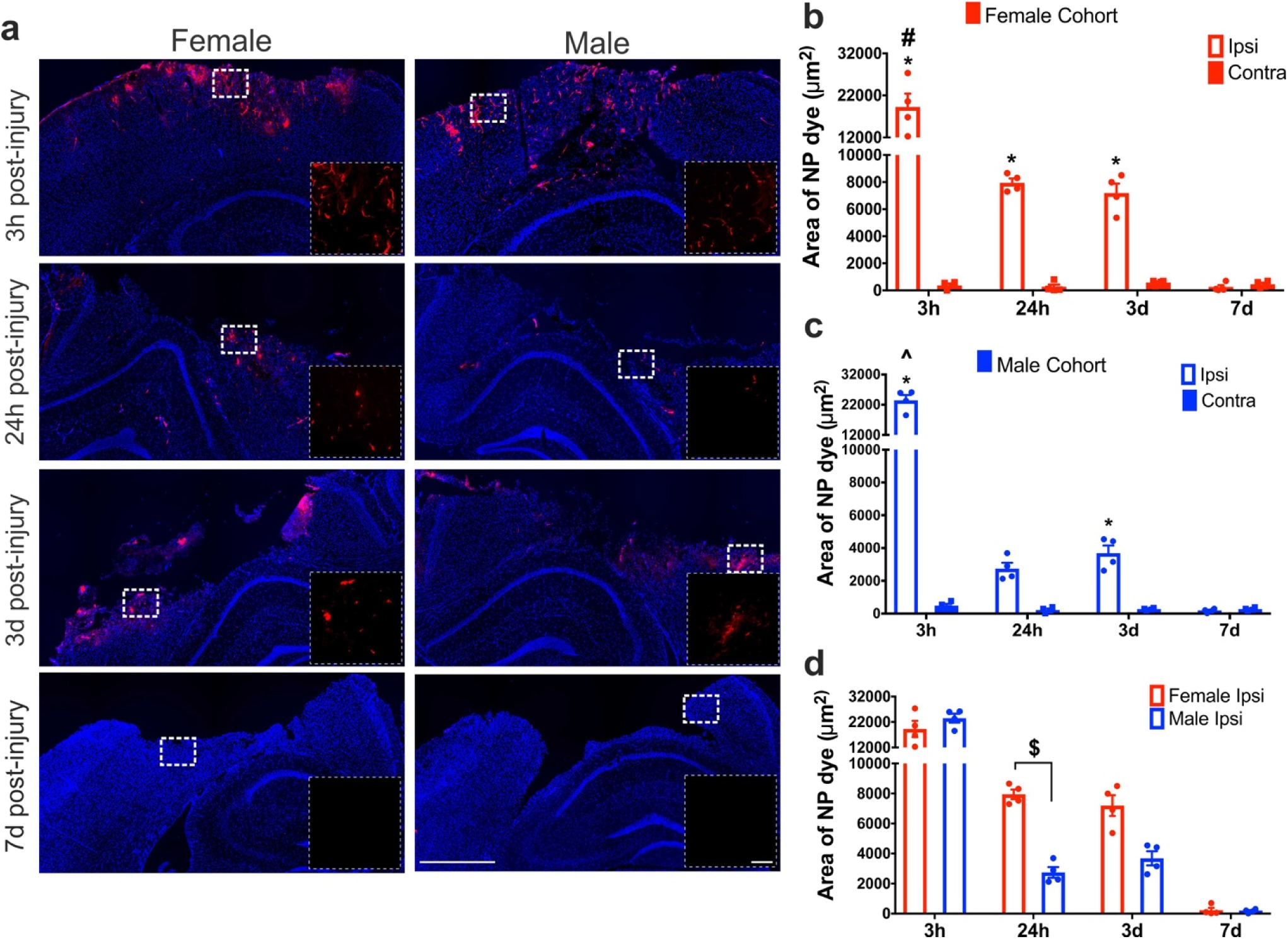
**(a)** NP accumulation after TBI in the ipsilateral hemisphere: Representative images of NP extravasation at 3 h, 24 h, 3 d and 7 days post-CCI. The first column shows the NP accumulation in the female cohort and the second column in males. The inset image is the enlarged view of the white dotted box in the main image. Main image scale bar = 750 μm, inset image scale bar = 100 μm. **(b-c)** NP accumulation in female and male cohort: Accumulation of NP in the female cohort (b) and male cohort (c) at different time points post-injury. *p<0.05 compared to their respective contralateral hemisphere and 7 d ipsilateral, two-way ANOVA, Bonferroni’s multiple comparisons. # p<0.05 compared to 24 h and 3 d ipsilateral hemisphere. ^ p<0.05 compared to 24 h ipsilateral hemisphere, $ p<0.05 compared to male 24 h ipsilateral hemisphere, two-way ANOVA, Bonferroni’s multiple comparisons. Error bars represent standard error of mean, n=4. **(d)** Quantification of NP accumulation in ipsilateral hemisphere of female and male cohorts across different time points.

Two-way ANOVA showed a significant main effect for both the hemisphere (p<0.0001) and time (p<0.0001) on HRP extravasation in both female (Fig. 2 (b)), and male cohorts (Fig. 2 (c)); the interaction between hemisphere and time was also significant (p<0.0001) (Supplementary Table 2 for statistical details). For the female cohort, the pair-wise analysis revealed significantly increased NP accumulation ipsilaterally at 3 h (p<0.0001), 24 h (p=0.0004) and 3 d (p=0.002) compared to their respective contralateral hemisphere. For the male cohort, a significant increase in NP accumulation was detected at 3 h (p <0.0001) and 3 d (p=0.0052) whereas the effect was marginal at 24 h (p=0.0508) post-injury compared to their respective contralateral location. At 7 d post-injury, both female (p>0.9999) and male cohorts (p>0.9999) did not show significant NP accumulation compared to their respective contralateral location. The 3 h ipsilateral NP accumulation was maximal for both female (p<0.0001, ∼150% increase) and male (p<0.0001, ∼500% increase) compared to all other time points (24 h, 3 d, and 7 d) in the ipsilateral hemisphere. Similar to HRP staining there was no significant difference in NP accumulation between 24 h and 3 d ipsilateral hemisphere in the female (p=0.9672) cohort. However, in contrast to the HRP staining, there was no significant difference in NP accumulation in males at 24 h and 3 d in the ipsilateral hemisphere (p=0.7509).

To investigate the sex dependence for NP delivery, we used two-way ANOVA to compare the female and male ipsilateral hemispheres across time points. Overall, the analysis revealed a significant interaction effect between sex and time (p=0.0087) with a significant main effect of time on NP accumulation (p<0.0001), but no main effect of sex on NP accumulation (p=0.2402) (Supplementary Table 2 for statistical details). A post-hoc pair-wise analysis at 24 h showed a significantly increased NP accumulation in females compared to males (p=0.0437), comparable to the HRP extravasation profile. Overall, we found significant NP accumulation in the ipsilateral hemisphere at acute and sub-acute time points in both female (3 h, 24 h, and 3 d) and male (3 h and 3 d) cohorts compared to the contralateral hemisphere. Furthermore, the ipsilateral hemisphere of the females showed significantly increased NP accumulation at 24 h post-injury compared to the male cohort.

### Intravital two photon microscopy imaging after intravenous injection of NP

The histological assessment demonstrated significant NP accumulation within the injury penumbra over the first three days after injury. However, tracking NP dynamics in post-mortem, perfused tissue sections is difficult, and artifacts related to perfusion-fixation itself may alter NP localization. Therefore, we employed intra-vital two-photon microscopic imaging for up to 3 h after NP injection for each post-injury cohort to decipher intravascular NP deposition versus extravasation into the parenchyma. The main advantage of two-photon microscopy imaging is the powerful resolution to visualize single cortical vessels and their surrounding cells after CCI. Moreover, we were able to image the dynamics of NP localization in the deep cortical regions (∼ 75 μm) post-injury in live tissue *in vivo*. (Supplementary videos and Fig 3). A combination of exogenous (via fluorescent NPs) and endogenous (via CX3CR1-GFP^+^) fluorescent labels were used to visualize the vascular and extravascular tissue. Transgenic mice with GFP tagged with CX3CR1 were used for direct visualization of microglia/macrophage cells present within the parenchyma, prominent inflammatory players followingTBI. A cranial window technique was used to enable imaging of GFP^+^ microglia cells and track the NPs with minimal perturbation to the normal physiological functions in live animals.

**Fig. 3:**
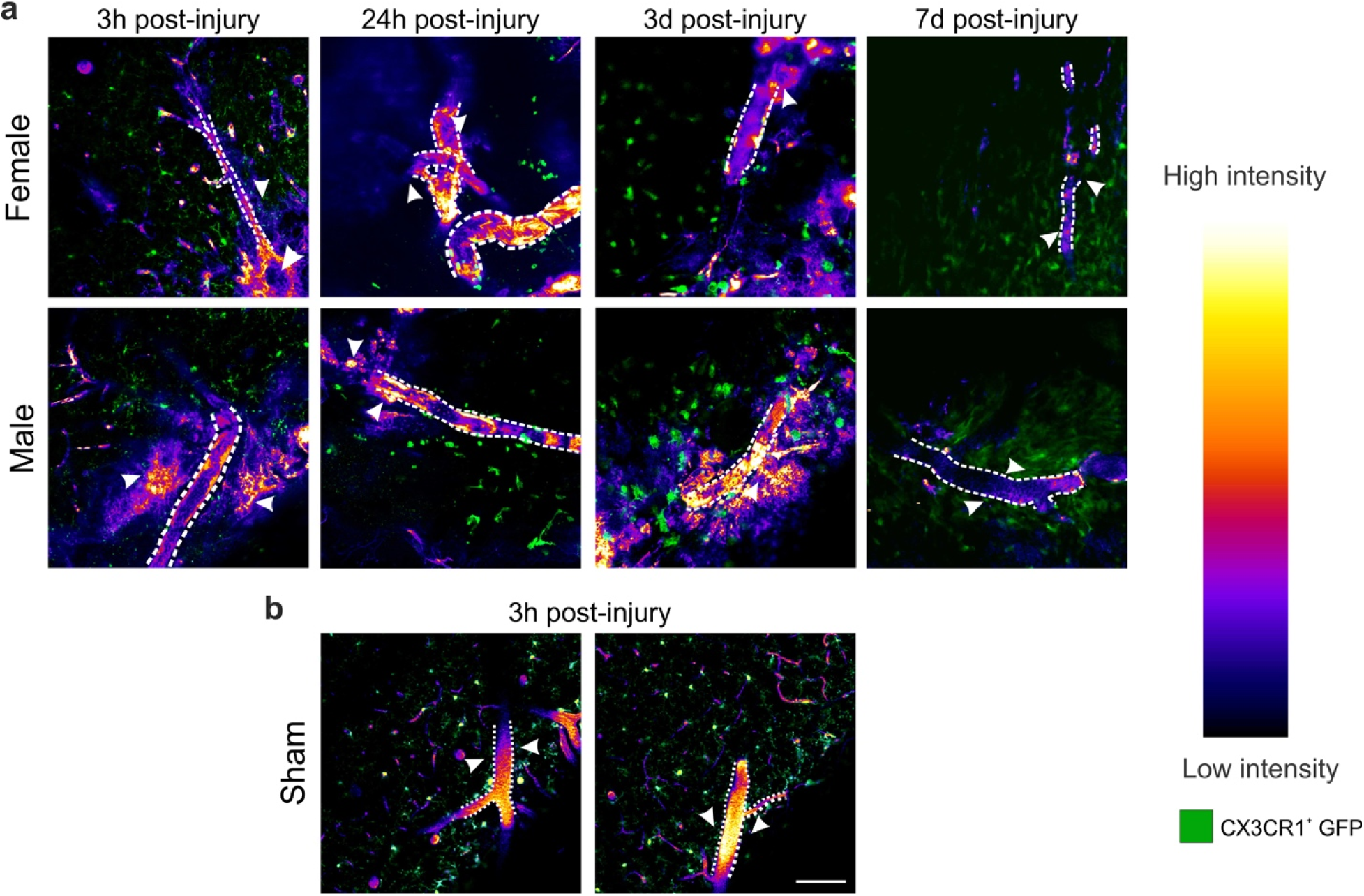
Two-photon intravital microscopy in the cerebral cortex after CCI. **(a)** Representative images of intravenously injected NPs (intensity map) acquired from the peri-injury zone of the cortex at 3 h, 24 h, 3 d, and 7 d post-CCI in CX3CR1-GFP^+^ transgenic mice (green) (blood vessels are denoted by a dotted white outline). Extravasation of NPs was evident in both male and female mice at 3 h, 24 h, and 3 d as diffuse labeling around the blood vessel (white arrows, shown here using a heatmap of grayscale intensity values on the right). At 7 d post-injury, there was no visible NP extravasation (white arrows around the blood vessels). **(b)** No extravasation or dispersed NP signal was observed in sham injury cohort animals (white arrows Fig. S6 shows the cranial window set up and the imaging location. Scale bar = 100 μm.

For each animal, a single two-photon video recording session was conducted for a maximum of 3 h after NP injection within the injury penumbra with as a minimum one scan per hour lasting at least 5 mins per scan. Overall, at least three different locations were imaged focusing on regions with identifiable, semi-intact blood vessels (See Supplementary Fig S6). Representative still images in Fig 3 (a), and supplementary videos show NP (in red) and the CX3CR1-GFP^+^ cells (microglia/macrophage; green). In the sham animals, we observed intact large blood vessels with the NP fluorescence and no NP extravasation, surrounded by ramified microglia (Fig 3 (b)). Additionally, we observed activated microglia with amoeboid morphology at 24 h and 3 d post-injury that corroborates previous studies^39–41^. We did not observe any detectable NP intensity within the extravascular tissue at 7 d post-injury in either sex.

### Injury lesion volume analysis: MRI analysis

MR images were used to quantify lesion volume size for sex comparison across the different time points (Fig 4). A longitudinal time series of 1 mm thick anatomically coronal T2-weighted spin-echo images (female, Fig 4a; male Fig 4b), show the evolution of the injury and lesion volume over time. It must be noted that the male cohort was imaged at 3 h, 24 h, 3 d, and 7 d; whereas the female cohort was imaged at 3h, 24 h, 4 d, and 7 d post-injury. The MR scans at 3 h and 24 h shows a typical signal hyperintensity at and around the injury site, characteristically interpreted as acute edema^42, 43^. By 3-4 d, this hyperintensity is reduced, suggesting the acute edema is resolving^42, 43^. At 7 d, there is typically an area of signal hyperintensity at the center of the injury region, usually interpreted as an area where tissue has been lost and replaced by fluid^42, 43^. Two-way ANOVA analysis showed no significant interaction between sex and time (p = 0.8084); yet, a significant main effect of both sex (p = 0.0017) and time (p = 0.0001) on lesion volume was identified (Supplementary Table 3 for statistical details). Pair-wise post-hoc analysis showed no significant differences between the female and male cohort for any time point. While these analyses indicate an overall main effect of sex on lesion size (i.e. females trended toward larger lesion volumes than males overall), within each time point no significant differences existed between sexes.

**Fig. 4.**
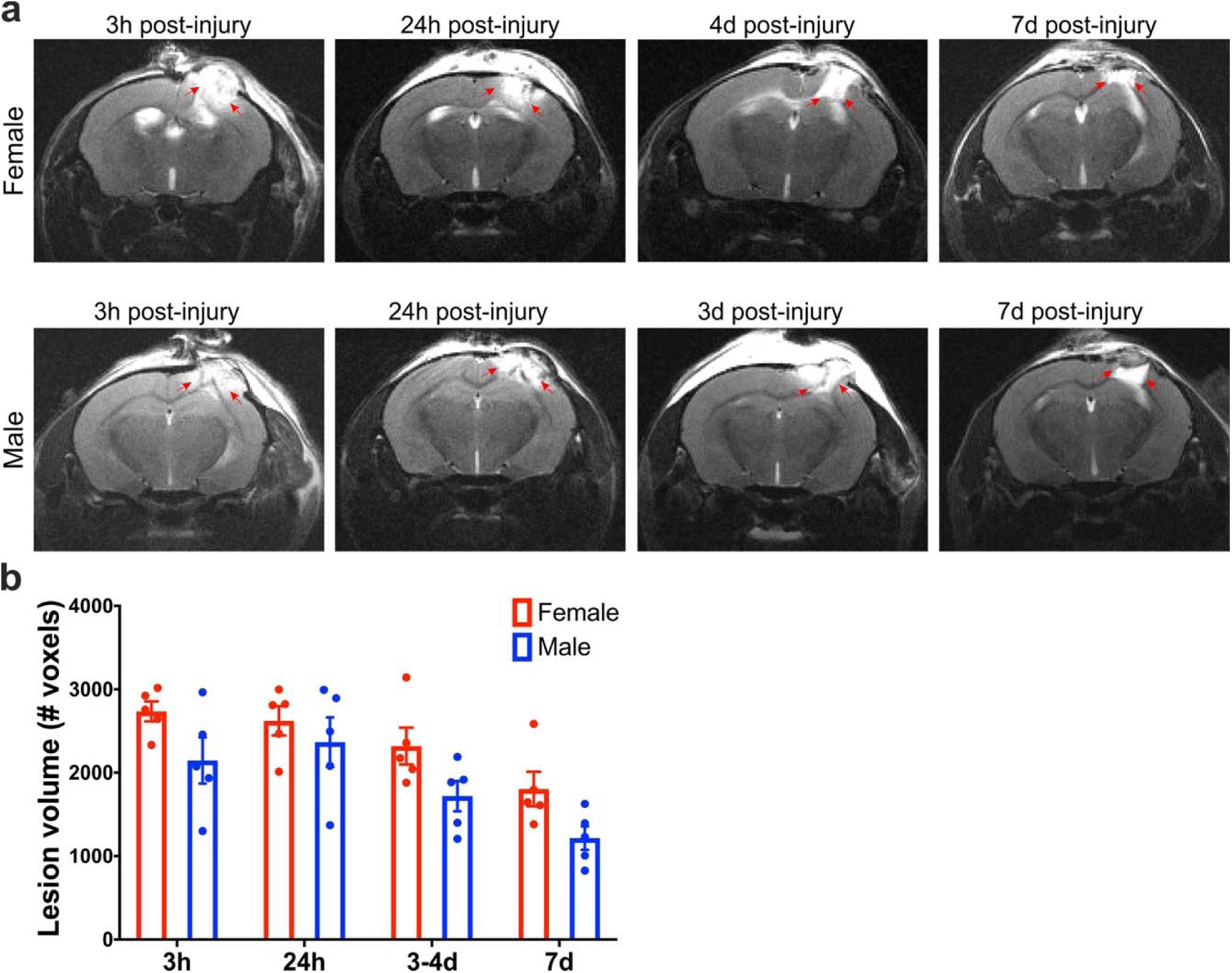
(a) Longitudinal in vivo T2 weighted magnetic resonance imaging (MRI) reveals changes in the cerebral cortex (right hemisphere) after CCI injury. A longitudinal cohort of female and male mice were repeatedly scanned at 3 h, 24 h, 3 d (male) or 4 d (female), and 7 d post-injury. Red arrows highlight the hyperintensity at and around the injury site. **(b)** Quantification of lesion volume T2 weighted scans acquired using a fast-spin echo sequence. Two-way ANOVA analysis, post-hoc Bonferroni’s analysis of the lesion volume comparison between sexes showed no significant difference at 3 h, 24 h, 3-4 d, and 7 d post-injury. Mean +/-SEM, n=5.

### Neuroglial response: Immunohistochemical analysis of glial fibrillary acidic protein (GFAP)

Neuroglial response in females and males in the ipsilateral and contralateral cortex was evaluated based on the level of reactive astrocytosis using glial fibrillary acidic protein (GFAP) immunostaining. The representative images of the ipsilateral hemisphere are shown in Fig 5 (a) and the contralateral images are shown in Supplementary Fig S7. Two-way ANOVA showed significant interaction between location (hemisphere) and time (p<0.0026, p<0.0002) and also a significant main effect for both hemisphere (p≤0.0002, p<0.0001) and time on GFAP immunostaining (p<0.0047, p<0.0001), respectively for female (Fig. 5 (b)), and male cohorts (Fig. 5 (c)) (Supplementary Table 4 for statistical details). We observed minimal GFAP expression at 3 h and 24 h post-injury for both female and male mice. For the female cohort, pair-wise analysis revealed significantly increased GFAP staining ipsilaterally at 3 d (p<0.0001) post-injury compared to the contralateral hemisphere. For the male cohort, a significant increase in GFAP staining was detected at 3 d (p <0.0001) and 7 d (p=0.0127) post-injury compared to their respective contralateral location. To investigate the sex dependence, a two-way ANOVA revealed a no significant interaction between sex and time nor a main effect of sex. Time was identified as a significant main effect on GFAP immunostaining levels (p<0.0001) (Fig 5 (d)). Taken together, the results show no significant sex difference in GFAP expression at any time point post-injury.

**Fig. 5.**
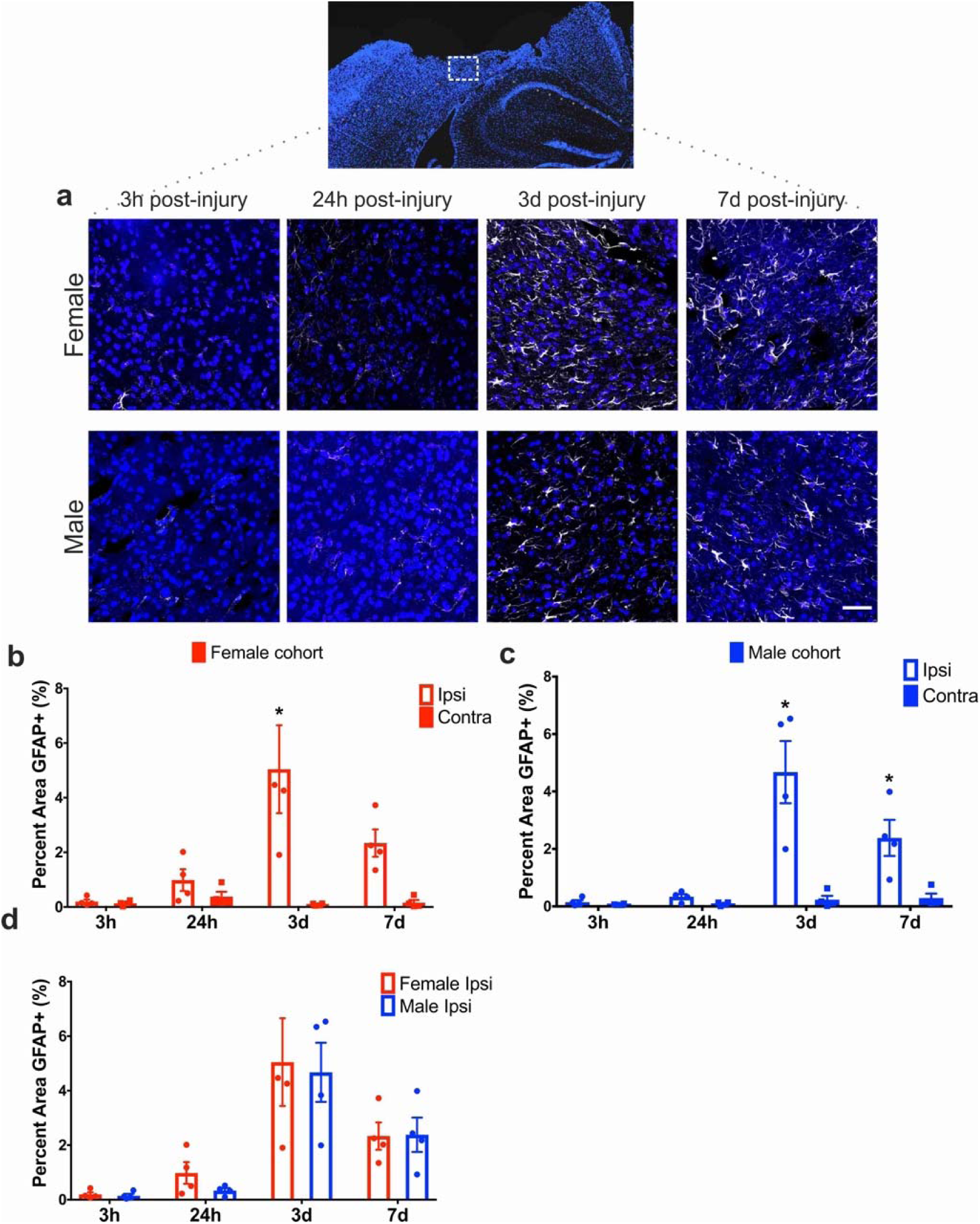
**(a)** Representative images of anti-glial fibrillary acidic protein (GFAP) staining within the cortical injury penumbra show progressive astrocyte reactivity at 3 h, 24 h, 3 d, and 7 d post-injury in both females and males. **(b)** Quantification of GFAP staining in the ipsilateral cortex found no difference between sexes at 3 h, 24 h, 3 d, and 7 d post-injury. Two-way ANOVA, Mean +/-SEM, n = 4 per group.

## Discussion

The blood-brain barrier (BBB) is a unique element of the brain that regulates molecular transport from the bloodstream to the brain parenchyma. Although the BBB disruption after TBI is thought to be detrimental, such opening may provide an opportunity for the delivery of drugs and therapeutics via NPs. To fully utilize the window of opportunity of BBB opening after TBI, a thorough assessment of the temporal resolution and the sex dependence of macromolecule for NP accumulation via the BBB was warranted. In this study, we directly address the critical gap using a focal TBI mouse model and intravenously injecting macromolecular tracer and PEGylated polystyrene NPs. The four key insights include: 1. Significant macromolecular tracer extravasation occurred at acute (3 h, 24 h) and sub-acute (3 d) post-injury time points after focal brain injury regardless of the sex; 2. Significant NP accumulation was observed acutely (3 h, 24 h in females and 3 h in males) and sub-acutely (3 d) after brain injury; 3. Sex-dependent effects were observed with a biphasic pattern in males showing a significant accumulation of the macromolecular tracer/NP at acute (3 h) and sub-acute (3 d) time point but a significant decrease at 24 h post-injury. In contrast, females showed a significant decrease at 24 h but continued to display significant accumulation within the first 3 d post-injury; 4. No sex differences were observed in the neuroglial response (GFAP) and while there was an overall main effect of sex on brain lesion volume (MRI analysis), no significant pairwise differences were observed at each time point.

Placing our data in context with prior studies that investigate differential sex responses to TBI, the following points are critical to consider. At our most acute (3 h) time point post-injury, we observed a peak accumulation of HRP and NP in both sexes with no significant sex differences. The controlled cortical impact model, consistently produces cortical damage including tearing of the dura, parenchyma and severe vascular disruption directly at the site of impact^29, 44^. Our findings of no sex differences 3 h post-injury in HRP and NP accumulation support mechanical disruption of the BBB from the primary impact. Comparing our data with previous studies using a diffuse TBI model, minimal BBB opening in intact (normal cycling) females was reported compared to significant BBB disruption in males at 5h post-injury^32, 33^. At 24 h post-injury, a sex-dependent BBB disruption emerged. Specifically, the females displayed a robust HRP/NP accumulation that was nearly 2.5-3x than their male counterparts. Although previous reports using different injury model^34^ and endogenous marker^35^ reported no sex differences in the BBB permeability at 24 h post-injury. The contrast between studies highlights the importance of considering injury phenotype (diffuse vs focal) and molecular tracer when comparing BBB disruption profiles. Notably, we designed our study to capture the dynamics of macromolecule and NP extravasation at distinct phases post-injury by intravenously injecting exogenous markers of NP (3 h) and HRP (10 mins) prior to sacrifice and perfusion at distinct time points post-injury. Moreover, the males showed a biphasic trend with a significant decrease in HRP/NP at 24 h compared to 3 h and 3 d post-injury, in alignment with previous seminal studies in male rodents using large molecular weight tracer^25, 45^. At sub-acute (3 d) time point, we observed significant macromolecular tracer and NP accumulation in both female and male cohorts compared to their respective contralateral region. Delayed BBB disruption is mainly associated with the secondary injury that follows the primary injury, although the mechanism(s) have not been clearly elucidated^9, 46^. Evidence suggests increased paracellular permeability due to disruption of tight junction complexes and integrity of the basement membrane as a dominant mechanism^9, 46, 47^. Oxidative stress due to reactive oxygen species (ROS) and free radical production after brain trauma alter the critical organization of tight junction proteins at the BBB resulting in increased paracellular leakage^9, 48^. Additionally, the disruption of vasculature and brain tissue caused by the primary impact triggers the coagulation cascade and leads to oxidative stress with increased production of proinflammatory mediators^9, 46^. These pathophysiological processes initiate glial cell activation and alter their interaction with the cerebrovascular endothelial cells potentially further contributing to BBB dysfunction^9, 46, 49, 50^. Taken together, to the best of our knowledge, this is the first report showing that the BBB disruption is sex-dependent for macromolecule and NP extravasation.

One of the key findings of our study is the differential sex response at 24 h post-injury between the macromolecular tracer (3 x higher) and NP accumulation (∼2.5 x higher) in females compared to males. These results were obtained from cohorts with no significant sex-dependent variations in astrogliosis and lesion volume. Numerous pre-clinical TBI studies report a female neuroprotective effect implicating estrogen or progesterone mediated mechanisms^58–61^, but a few studies report no impact of sex on the outcome^62, 63^. In contrast, clinical studies cite worse outcomes for females compared to males^64–68^. Our study design mirrored prior work that indicated that the phase of the estrous cycle at the time of injury does not significantly affect functional outcomes following TBI^69^. Yet, future studies are warranted as a study by Maghool et. al. in rats reported females in proestrus phase are less vulnerable to brain injury compared with females in non-proestrus phase as measured by levels of brain edema and intracranial pressure^70^. Furthermore, it is important to recognize that a TBI event may alter subsequent sex hormone levels thereby affecting estrous cycles acutely after injury. A pre-clinical rat study shows evidence that TBI leads to disruption of the estrous cycle with significantly reduced 17-estradiol (E2) hormone level^71^ and spatial memory impairment compared to the sham group. Similarly, clinical studies indicate that TBI affects the circulating serum hormone levels in both females and males^72, 73^. In females, testosterone increased with modest/no increase in estrogens^72^. In males, estrogen increases with a decrease in testosterone^73^. Therefore, specific studies to understand the role of sex hormones in neuroprotection and BBB breakdown after TBI are warranted.

In conclusion, to the best of our knowledge, we report for the first time that (1) female mice exhibit a robust macromolecular tracer (3 x higher) and NP accumulation (2.5 x higher) at 24 h time point after focal TBI compared to male mice, primarily due to sex differences in BBB permeability in response to the injury and (2) we demonstrate the ability to delivery macromolecules and NPs at sub-acute time point (3 d) after focal injury in both female and male mice. The implications of our novel findings are far-reaching for personalized and precision medicine. The sex differences in macromolecular and NP extravasation may impact the pharmacokinetics and pharmacodynamics of the therapeutic delivery after brain injury. Moreover, the sex-based differences in extravasation may influence the drug dose, drug concentration and desired bioavailability near the injured brain tissue between female and male subjects. Therefore, a better understanding of the sex differences is essential to appropriately conduct a risk assessment and to design safe and effective treatments. Future studies to elucidate the underlying hormonal and sex-related differences for variable BBB permeability are warranted.

## Materials and methods

### Materials

Carboxylated polystyrene NPs of 40 nm (F8793) size with red (λ_ex_/λ_em_ = 580 nm/605 nm) was purchased from Life Technologies (Carlsbad, CA, USA). Methoxypolyethylene glycol amine 2000 (mPEGamine 2 KDa) (06676), n-[3-dimethylaminopropyl]-n-ethyl, n-[3-dimethylaminopropyl]-n-ethyl [EDC] (E1769), MES hemisodium buffer (M8902), N-Hydroxysuccinimide (NHS) (56405), and Peroxidase type II from horseradish (P8250-50KU) were purchased from Sigma Aldrich (St. Louis, MO, USA). ImmPACT DAB peroxidase (HRP) substrate (SK-4105) was purchased from Vector laboratories (Burlingame, CA, USA). Slide-A-Lyzer Cassettes (20 K) (66003) were purchased from ThermoFisher scientific (Waltham, MA, USA). Vectashield antifade mounting medium (H-1000) from Vector Labs (Burlingame, Ca, USA) were purchased. Anti-GFAP (ab53554) was purchased from Abcam (Cambridge, MA, USA). Alexa Fluor 647 Secondary (ab150131) was purchased from Abcam (Cambridge, MA, USA).

### Nanoparticle PEG conjugation

As presented in our previous study^19, 26^, carboxylated NPs were PEGylated using EDC/NHS chemistry. Briefly, mPEGamine 2 kDa was mixed with 40 nm (NH_2_: COOH at 5:1 mole excess). EDC/NHS (in MES buffer) was added to NP / PEG mixture (200 mM/100 mM) and HEPES buffer was added to obtain a final pH of 7.8 before incubating for 3 h at room temperature. Glycine (100 mM) was added to quench the reaction. Unbound PEG was removed via dialysis (20 kDa MW). PEGylated NPs were suspended in a 20 mM HEPES (pH 7.4). The concentration of NP solution was determined with fluorescent standard curves generated from known concentrations of as-received Fluorospheres (FLUOstar Omega fluorescence plate reader; BMG Labtech, Ortenberg, Germany). Yields of NPs ranged between 40-60 %. A concentration of 12.5 mg/mL for each NP was used for all *in vivo* studies.

### Nanoparticle characterization

The size (in nanometers) and surface charge (zeta potential in millivolts) of the pre- and post-PEGylation of NPs were evaluated using a Zeta Sizer Nano-ZS (Malvern Instruments, Malvern, UK). Each NP sample in 20 mM HEPES (pH 7.4) was measured three times consecutively. Three measurements were made and the mean ± standard deviation (s.d.) was reported. To study the stability of the NP for 3 h post-injection, unPEGylated NPs and PEGylated NPs were diluted in serum (incubated at 37°C and 5% CO_2_ for at least 2h before use) at a concentration comparable to the *in vivo* blood concentration (0.42 mg/ml). The samples were then incubated in a 96-well plate at 37°C and 5% CO_2_ incubator for a about 3 h and the aggregation was monitored by measurement of absorbance at 320 nm at 37°C.

### NMR characterization

The conjugation of PEG and the total quantity on NP was detected by ^1^H NMR spectra analysis using a previously published methodology^74^. PEGylated and unPEGylated polystyrene NPs were weighed and fully dissolved in a mixture of chloroform (CDCl_3_, Sigma), trifluoroacetic acid-d (TFAd, Sigma), and a known concentration BTSB (0.5% w/v). ^1^H NMR spectra were obtained at 400 MHz using a Varian Inc. (Palo Alto, CA, USA) VNMRS 500 MHz NMR spectrometer with a 5 mm ID-PFG probe. PEG2000 polymer (3.6 ppm) was dissolved in the same CDCL_3_-TFAd solvent with BTSB (0.5% w/v) was made at different concentrations to obtain a calibration curve. Total PEG quantity in the PEGylated NP was calculated using the calibration curve.

### Animals

Female and male adult C57BL/6 mice (Jackson Laboratory) aged 8-10 weeks (20-24 g) were used for the HRP extravasation, NP accumulation, IHC analysis experiments (n=4 per group). Female and male adult transgenic CX3CR1-GFP (Jackson Laboratory) mice bred with C57BL/6 mice (Jackson Laboratory) were used for two-photon microscopy experiments (n = 3 per group). Another second cohort of female and male adult C57BL/6 mice aged 8-10 weeks (20-24 g) were used for MRI lesion volume analysis and biodistribution and blood plasma concentration analysis (n = 5). Mice were housed in a 14 h light/10 h dark cycle at a constant temperature (23°C ± 2° C) with food and water available *ad libitum*. Female mice of random cycling were used. Animal studies using C57BL/6 mice were approved by Arizona State University’s Institutional Animal Use and Care Committee (IACUC) and were performed in accordance with the relevant guidelines. The two-photon microscopy study using transgenic animals was approved by the Institutional Animal Care and Use Committee at the University of Arizona (Tucson, AZ).

### Controlled cortical impact model

Traumatic brain injury (TBI) was modeled using the well-established controlled cortical impact (CCI) injury model^75^. Briefly, anesthetized (isoflurane) adult mice were mounted onto a stereotaxic frame. The frontoparietal cortex was exposed via 3-4 mm craniotomy and the impact tip was centered to the craniectomy. The impactor tip diameter was 2 mm, the impact velocity was 6.0 m/s and the depth of cortical deformation was 2 mm and 100 ms impact duration (Impact ONE; Leica Microsystems). The skin was sutured and the animals were placed in a 37°C incubator until consciousness was regained. The sham group went through the same protocol but did not receive the impact injury.

### Nanoparticle (NP) and horseradish peroxidase (HRP) injection

Retro-orbital injections of the venous sinus in the mouse, an alternative technique to tail-vein injection^76^, were performed for intravenous delivery of the NPs and HRP. Animals were anesthetized with isoflurane (3 %) and received intravenous NP injections (30 mg/kg b.w. 50 μL; retro-orbital) of 40 nm NPs three hours before perfusion. HRP (83 mg/kg b.w.in 25 μL) was retro-orbitally injected behind the second eye ten minutes before perfusion. Depending on the cohort group, animals were sacrificed at 3 h, 24 h, 3 d or 7 d post-injury. The NP circulation time of 3 h was held constant for all cohorts. The summary of the experimental timeline is depicted in Supplementary figure 1.

### Tissue collection

At the end time point, mice were deeply anesthetized with a lethal dose of sodium pentobarbital solution. After the loss of a tail/toe pinch reflex movement, animals were transcardially perfused with cold phosphate-buffered saline (PBS), followed by 4 % buffered paraformaldehyde solution. Brain tissue was collected and fixed overnight in 4 % (w/v) buffered paraformaldehyde followed by immersion in 30 % (w/v) sucrose solutions in 1X PBS for cryoprotection until the tissue was fully infiltrated. Samples were embedded in optimal cutting temperature (OCT) medium and frozen by placing in a glass container with methylbutane kept on dry ice. Samples were stored at −80°C until sectioned coronally at a 20 μm thickness with a cryostat (Cryostar^TM^ NX70, ThermoFisher scientific (Waltham, MA, USA)).

### Quantification of HRP extravasation

The tissue section was incubated in PBS buffer for 20 mins at room temperature prior to use. The tissue sampling regions were ∼-1.65 mm Bregma (four sections per animal, n=4 animals per group). For HRP analysis, freshly prepared DAB substrate solution (200 μL) was added to the tissue and incubated for ten mins at room temperature. Slides were then washed in deionized water three times (two mins each) and coverslips were mounted after adding a drop of aqueous mounting media. Sections were imaged using Slide Scanner (PathScan Enabler IV, Meyer Instruments, TX, USA). Region of interest of dimension 5.2 mm x 2.5 mm was selected around the injury penumbra including the cortical and the hippocampus regions. The ROI images were then analyzed using ImageJ software (National Institute of Health, Bethesda, MD, USA) to obtain the total number of positive pixels per ROI.

### Quantification of NP accumulation

For NP analysis, similar to HRP analysis, slides containing the frozen sections were incubated at room temperature for 20 mins in 1X PBS to rehydrate the tissue and remove OCT compound. Coverslips were mounted on the section after adding one drop of fluorescent mounting media (Vectashield). The tissue sampling regions were ∼-1.65 mm Bregma and ipsilateral cortex (four sections per animal; n = 4) and contralateral cortex (two sections per animal, n = 4) were imaged with conventional epi-fluorescent microscopy at 10 X objective. All images were taken at the same exposure time and consistent acquisition settings. Similar to HRP analysis, a region of interest 5.2 mm x 2.5 mm was selected around the injury penumbra including the cortical and the hippocampus regions. The ROI images were then analyzed using ImageJ software (National Institute of Health, Bethesda, MD, USA). A threshold was set using images from sham cohort tissues to remove background noise and was kept constant for ipsilateral and contralateral image analysis. The thresholded images were used to obtain the total number of positive pixels per ROI.

### Two-photon microscopy: Cranial window placement and Imaging

Transgenic mice (female and male) underwent the CCI injury induction as described above. Based on the imaging time points (immediately after the injury or at 24 h, 3 d, or 7 d post-injury), anesthetized animals received intravenous NP injections (30 mg/kg b.w., 50 μL; retro-orbital). Following the injection, a cranial window was placed for imaging. The cranial window protocol is modified from a previously published protocol^77, 78^. Briefly, the craniectomy/injury region was cleaned using saline and gel foam soaked in saline was applied to stop any bleeding (if any). The region around the craniectomy was dried using a cotton swab. After ensuring no bleeding, 5 mm diameter glass coverslip was gently placed on the brain tissue, centered at −2 mm bregma and 2 mm lateral midline, Fig. S6 (a-b). The glass coverslip was sealed using dental cement to secure it and to create a well for the water-immersion objective. One cohort of control, sham animals were used to establish baseline NP extravasation. These sham animals were anesthetized, injected with NPs, then subjected to craniectomy (4 mm), and cranial window placement according to the aforementioned protocol. For imaging, animals remained under anesthesia (isoflurane) and were mounted on a microscope adaptable animal holding frame. Animals were placed on the mount using ear bars to stabilize their heads and two-photon laser scanning microscope (FVMPE-RS) from Olympus (Tokyo, Japan) was used. A water immersion 25 X objective lens (NA = 1.05) was used to image NPs present inside the cerebral microvessels and diffuse NP signal indicating NP extravasation outside of the blood vessels. 12-bit images of 512 × 512 pixels (0.509 × 0.509 mm in the x-y plane) were obtained with a Galvano scanner via one-way scan direction and sampling speed of 4.0 μs/pixel. The excitation wavelength of 920 nm was used for the red channel and green channel with ∼15 % laser transmissivity. Imaging was completed at a depth ∼75-80 μm below from the coverslip to avoid any artifact related to superficial bleeds that arise during coverslip placement. Animals were imaged for a maximum of 3 h under anesthesia (isoflurane). After two-photon imaging was completed, while animals were still under anesthesia, horseradish peroxidase (83 mg/kg of b.w. injection volume of 25 μL) was injected via retro-orbital 10 mins before sacrifice. Animals were perfused using the protocol explained above.

### MRI lesion volume analysis

MRI was performed longitudinally at 3 h, 24 h, 3-4 d (3 d for male cohort and 4 d for female cohort) and 7 d post-injury using ParaVision software on a Bruker Biospin 7T system (Bruker Biospin Corp., Billerica, MA). A volume transmitter coil (72 mm) was used in conjunction with a mouse brain surface coil for signal detection. Animals were placed at a prone position on a nonmagnetic holder with the teeth bar as an aid to fix the head position. During image acquisition, anesthesia was maintained using isoflurane (1.5 %) while the body temperature was maintained at 37°C using a circulating water blanket. Respiration and rectal temperature were monitored using SAI system. T2 weighted scans were acquired using a fast-spin echo sequence (TR/TE/ETL= 4000 ms /60 ms /8, resolution of 0.1 mm X 0.1 mm X 0.5 mm, 30 slices). The T2-weighted scans were then imported into ImageJ for lesion volume analysis. For each scan, freehand ROIs were drawn around the injury for each slice. The ROI/injury was determined by visually evaluating the area of signal hyperintensity around the location where the cortical impact occurred. The lesion volume was computed for a scan by summing the area of the ROIs of each slice of that scan in terms of the number of voxels. This volume was found for each animal for each time point.

### Immunohistochemical staining, imaging and quantification

Four tissue sections per animal were analyzed. Tissue sections were washed four times for 15 minutes between all stains. Initially, sections were blocked for one hour using 8 % horse serum and 0.2 % TritonX in 1X PBS. Sections were incubated overnight at 4°C in 1:500 goat anti-GFAP, 2 % horse serum and 0.1 % Triton X in 1X PBS. Slides were then incubated for two hours in 1:500 donkey anti-goat 647 secondary with 2 % horse serum in 1X PBS in dark. Finally, slides were stained with DAPI, mounted with vectashield and sealed with nail polish. Slides were imaged at 20 X on the Zeiss LSM800 Laser scanning confocal microscope. The region of interest for imaging was selected at the penumbra region of the injury site, ∼200-500 μm away from the lesion site to avoid the necrosis region. DAPI and GFAP image acquisition settings were held constant. Images were taken around the injury and on the contralateral hemisphere. The percent area was calculated using ImageJ. A threshold was set using tissues from secondary control images to remove background noise and was kept constant for ipsilateral and contralateral images. The thresholded images (ipsilateral and contralateral) were quantified using analyze particles. The percent area result was then averaged per animal.

### Statistics

Statistical analyses were conducted in GraphPad Prism 5.0 (GraphPad Software, Inc., La Jolla CA). For HRP, NP, and GFAP analysis, an ordinary two-way ANOVA was conducted to analyze female and male cohorts individually. If there was a significant main effect of hemisphere (ipsilateral vs contralateral) and/or significant interaction effect, the two-way ANOVA analysis was followed by a post-hoc Bonferroni’s multiple comparison tests. To evaluate the sex dependence for HRP, NP, and GFAP analysis, an ordinary two-way ANOVA was conducted using ipsilateral hemispheres of female and male cohorts. If there was a significant main effect of sex (female vs male) and/or significant interaction effect, the two-way ANOVA analysis was followed by a post-hoc Bonferroni’s multiple comparison tests. For MRI analysis, a two-way ANOVA with replication was conducted to analyze the effect of both time and sex on injury lesion volume. If there was a significant main effect of sex (female vs male) and/or significant interaction effect, the two-way ANOVA analysis was followed by a post-hoc Bonferroni’s multiple comparison tests.

## Supporting information

Two-Photon Videos Female Cohort

Two-Photon Videos Male Cohort

Sham Cohort

## Acknowledgements

Authors would like to thank Katherine R. Giordano for assisting in breeding the transgenic animals and Chen Wu for helping with perfusions of the transgenic animals. We thank Dr. Shenfeng Qiu for lending the animal mount for two-photon microscope imaging study. We thank Jordan Todd and Kyle Offenbacher for their technical assistance. We thank the Barrow-Arizona State University Center for Preclinical Imaging for the use of magnetic resonance imaging and IVIS spectrum. We thank Xiaowei Zhang for performing the MRI imaging. We thank Dr. Racheal Sirianni for the use of dynamic light scattering device. Authors thank Brian Cherry and Magnetic Resonance Research Center at Arizona State University for NMR analysis. We thank the Keck imaging facility for the fluorescent spectrophotometer use. This study was supported by the FLINN Foundation (S.E.S., V.D.K., J.L., P.D.A.), NIH (R00NS076661 and R01NS097537 to JMN; 1DP2HD084067 to SES) and Arizona State University Graduate College Completion Fellowship (V.N.B.).

## References

1. Pardridge, W. M. The blood-brain barrier: bottleneck in brain drug development. Neurotherapeutics 2, 3–14 (2005).

2. de Boer, A. G. & Gaillard, P. J. Drug targeting to the brain. Annu. Rev. Pharmacol. Toxicol. 47, 323–355 (2007).

3. Pardridge, W. M. Drug transport across the blood-brain barrier. Journal of Cerebral Blood Flow & Metabolism 32, 1959–1972 (2012).

4. Banks, W. A. From blood–brain barrier to blood– brain interface: new opportunities for CNS drug delivery. Nature Publishing Group 15, 275–292 (2016).

5. Saunders, N. R., Habgood, M. D., Møllgård, K. & Dziegielewska, K. M. The biological significance of brain barrier mechanisms: help or hindrance in drug delivery to the central nervous system? F1000Res 5, 313–15 (2016).

6. Davis, A. E. Mechanisms of traumatic brain injury: biomechanical, structural and cellular considerations. Crit Care Nurs Q 23, 1–13 (2000).

7. Gennarelli, T. A. The Pathobiology of Traumatic Brain Injury. Neuroscientist 3, 73–81 (1997).

8. Baldwin, S. A., Fugaccia, I., Brown, D. R., Brown, L. V. & Scheff, S. W. Blood-brain barrier breach following cortical contusion in the rat. Journal of neurosurgery 85, 476–481 (1996).

9. Shlosberg, D., Benifla, M., Kaufer, D. & Friedman, A. Blood-brain barrier breakdown as a therapeutic target in traumatic brain injury. 6, 393–403 (2010).

10. Clond, M. A. et al. Reactive oxygen species-activated nanoprodrug of Ibuprofen for targeting traumatic brain injury in mice. PLoS ONE 8, e61819–e61819 (2013).

11. Bharadwaj, V. N., Nguyen, D. T., Kodibagkar, V. D. & Stabenfeldt, S. E. Nanoparticle-Based Therapeutics for Brain Injury. Adv. Healthcare Mater. 7, 1700668–16 (2017).

12. Dufès, C. Brain Delivery of Peptides and Proteins. Peptide and Protein Delivery 105–122 (Elsevier, 2011). doi:10.1016/B978-0-12-384935-9.10006-9

13. Olivier, J.-C. Drug transport to brain with targeted nanoparticles. Neurotherapeutics 2, 108–119 (2005).

14. Agarwal, A. et al. Nanoparticles as novel carrier for brain delivery: a review. Curr. Pharm. Des. 15, 917–925 (2009).

15. Petros, R. A. & DeSimone, J. M. Strategies in the design of nanoparticles for therapeutic applications. Nat Rev Drug Discov 9, 615–627 (2010).

16. Parveen, S., Misra, R. & Sahoo, S. K. Nanoparticles: a boon to drug delivery, therapeutics, diagnostics and imaging. Nanomedicine: Nanotechnology, Biology, and Medicine 8, 147–166 (2012).

17. Masserini, M. Nanoparticles for Brain Drug Delivery. ISRN Biochem 2013, 1–18 (2013).

18. Meyers, J. D., Doane, T., Burda, C. & Basilion, J. P. Nanoparticles for imaging and treating brain cancer. Nanomedicine 8, 123–143 (2013).

19. Bharadwaj, V. N. et al. Blood-brainbarrier disruption dictates nanoparticle accumulation following experimental brain injury. Nanomedicine: Nanotechnology, Biology, and Medicine 14, 2155–2166 (2018).

20. Fosgerau, K. & Hoffmann, T. Peptide therapeutics: current status and future directions. Drug Discov. Today 20, 122–128 (2015).

21. Usmani, S. S. et al. THPdb: Database of FDA-approved peptide and protein therapeutics. PLoS ONE 12, e0181748 (2017).

22. Bobo, D., Robinson, K. J., Islam, J., Thurecht, K. J. & Corrie, S. R. Nanoparticle-Based Medicines: A Review of FDA-Approved Materials and Clinical Trials to Date. Current Opinion in Solid State and Materials Science 33, 2373–2387 (2016).

23. Ventola, C. L. Progress in Nanomedicine: Approved and Investigational Nanodrugs. P T 42, 742–755 (2017).

24. Habgood, M. D. et al. Changes in blood-brain barrier permeability to large and small molecules following traumatic brain injury in mice. Eur. J. Neurosci. 25, 231–238 (2007).

25. Baskaya, M. K., Muralikrishna Rao, A., Dogan, A., Donaldson, D. & Dempsey, R. J. The biphasic opening of the blood–brain barrier in the cortex and hippocampus after traumatic brain injury in rats. Neurosci. Lett. 226, 33–36 (1997).

26. Bharadwaj, V. N., Lifshitz, J., Adelson, P. D., Kodibagkar, V. D. & Stabenfeldt, S. E. Temporal assessment of nanoparticle accumulation after experimental brain injury: Effect of particle size. Sci. Rep. 6, 29988 (2016).

27. Ping, X., Jiang, K., Lee, S.-Y., Cheng, J.-X. & Jin, X. PEG-PDLLA micelle treatment improves axonal function of the corpus callosum following traumatic brain injury. J Neurotrauma 31, 1172–1179 (2014).

28. Bharadwaj, V. N., Nguyen, D. T., Kodibagkar, V. D. & Stabenfeldt, S. E. Nanoparticle-Based Therapeutics for Brain Injury. Adv. Healthcare Mater. 7, (2018).

29. Baldwin, S. A., Fugaccia, I., Brown, D. R., Brown, L. V. & Scheff, S. W. Blood-brain barrier breach following cortical contusion in the rat. Journal of neurosurgery 85, 476–481 (1996).

30. Vagnerova, K., Koerner, I. P. & Hurn, P. D. Gender and the Injured Brain. Anesthesia & Analgesia 107, 201–214 (2008).

31. Caplan, H. W., Cox, C. S. & Bedi, S. S. Do microglia play a role in sex differences in TBI? Journal of Neuroscience Research 95, 509–517 (2017).

32. O’Connor, C. A., Cernak, I. & Vink, R. Both estrogen and progesterone attenuate edema formation following diffuse traumatic brain injury in rats. Brain Research 1062, 171–174 (2005).

33. O’Connor, C. A., Cernak, I. & Vink, R. in Brain Edema XIII 96, 121–124 (Springer, Vienna, 2006).

34. Duvdevani, R., Roof, R. L., Fulop, Z., Hoffman, S. W. & Stein, D. G. Blood-brain barrier breakdown and edema formation following frontal cortical contusion: does hormonal status play a role? http://www.liebertpub.com/neu 12, 65–75 (1995).

35. Jullienne, A. et al. Male and Female Mice Exhibit Divergent Responses of the Cortical Vasculature to Traumatic Brain Injury. http://www.liebertpub.com/neu 35, 1646–1658 (2018).

36. Stewart, P. A., Farrell, C. R., Farrell, C. L. & Hayakawa, E. Horseradish peroxidase retention and washout in blood-brain barrier lesions. Journal of Neuroscience Methods 41, 75–84 (1992).

37. Tanno, H., Nockels, R. P., Pitts, L. H. & Noble, L. J. Breakdown of the blood-brain barrier after fluid percussive brain injury in the rat. Part 1: Distribution and time course of protein extravasation. J Neurotrauma 9, 21–32 (1992).

38. Lotocki, G. et al. Alterations in blood-brain barrier permeability to large and small molecules and leukocyte accumulation after traumatic brain injury: effects of post-traumatic hypothermia. J Neurotrauma 26, 1123–1134 (2009).

39. Chen, S. Time course of cellular pathology after controlled cortical impact injury. Experimental Neurology 182, 87–102 (2003).

40. Turtzo, L. et al. Macrophagic and microglial responses after focal traumatic brain injury in the female rat. Journal of Neuroinflammation 11, 82–14 (2014).

41. Villapol, S., Loane, D. J. & Burns, M. P. Sexual dimorphism in the inflammatory response to traumatic brain injury. Glia 65, 1423–1438 (2017).

42. Kochanek, P. M. et al. Severe controlled cortical impact in rats: assessment of cerebral edema, blood flow, and contusion volume. http://www.liebertpub.com/neu 12, 1015–1025 (1995).

43. Onyszchuk, G. et al. A mouse model of sensorimotor controlled cortical impact: Characterization using longitudinal magnetic resonance imaging, behavioral assessments and histology. Journal of Neuroscience Methods 160, 187–196 (2007).

44. Adelson, P. D., Whalen, M. J., Kochanek, P. M., Robichaud, P. & Carlos, T. M. Blood brain barrier permeability and acute inflammation in two models of traumatic brain injury in the immature rat: a preliminary report. 71, 104–106 (1998).

45. Huang, Z. G., Xue, D., Preston, E., Karbalai, H. & Buchan, A. M. Biphasic opening of the blood-brain barrier following transient focal ischemia: effects of hypothermia. Can J Neurol Sci 26, 298–304 (1999).

46. Chodobski, A., Zink, B. J. & Szmydynger-Chodobska, J. Blood-brain barrier pathophysiology in traumatic brain injury. Transl. Stroke Res. 2, 492–516 (2011).

47. Sandoval, K. E. & Witt, K. A. Blood-brain barrier tight junction permeability and ischemic stroke. Neurobiology of Disease 32, 200–219 (2008).

48. Pardridge, W. M. Blood-brain barrier biology and methodology. J. Neurovirol. 5, 556–569 (1999).

49. Chen, Y. & Swanson, R. A. Astrocytes and Brain Injury. J. Cereb. Blood Flow Metab. 23, 137–149 (2003).

50. Karve, I. P., Taylor, J. M. & Crack, P. J. The contribution of astrocytes and microglia to traumatic brain injury. British journal of pharmacology (2016). doi:10.1111/bph.2016.173.issue-4

51. Hall, E. D., Gibson, T. R. & Pavel, K. M. Lack of a gender difference in post-traumatic neurodegeneration in the mouse controlled cortical impact injury model. http://www.liebertpub.com/neu 22, 669–679 (2005).

52. Tucker, L. B., Fu, A. H. & McCabe, J. T. Performance of Male and Female C57BL/6J Mice on Motor and Cognitive Tasks Commonly Used in Pre-Clinical Traumatic Brain Injury Research. http://www.liebertpub.com/neu 33, 880–894 (2016).

53. Tucker, L. B., Burke, J. F., Fu, A. H. & McCabe, J. T. Neuropsychiatric Symptom Modeling in Male and Female C57BL/6J Mice after Experimental Traumatic Brain Injury. http://www.liebertpub.com/neu 34, 890–905 (2017).

54. Bruce-Keller, A. J. et al. Gender and estrogen manipulation do not affect traumatic brain injury in mice. http://www.liebertpub.com/neu 24, 203–215 (2007).

55. Fonseca, E. A., Duran, J. C., Carrero, P., Segura, L. M. G. & Arevalo, M. A. Sex differences in glia reactivity after cortical brain injury. Glia 63, 1966–1981 (2015).

56. Günther, M. et al. COX-2 regulation and TUNEL-positive cell death differ between genders in the secondary inflammatory response following experimental penetrating focal brain injury in rats. Acta Neurochir (Wien) 157, 649–659 (2015).

57. Clevenger, A. C. et al. Endogenous Sex Steroids Dampen Neuroinflammation and Improve Outcome of Traumatic Brain Injury in Mice. 1–11 (2018). doi:10.1007/s12031-018-1038-x

58. Roof, R. L. & Hall, E. Gender differences in acute CNS trauma and stroke: neuroprotective effects of estrogen and progesterone. Journal of Nuerotrauma 17, (2000).

59. Bramlett, H. M. & Dietrich, W. D. Neuropathological Protection after Traumatic Brain Injury in Intact Female Rats Versus Males or Ovariectomized Females. http://www.liebertpub.com/neu 18, 891–900 (2004).

60. Stein, D. G. Brain damage, sex hormones and recovery: a new role for progesterone and estrogen? Trends in Neurosciences 24, 386–391 (2001).

61. Stein, D. G. Brain Trauma, Sex Hormones, Neuronal Survival, and Recovery of Function. Principles of Gender-Specific Medicine 104–115 (Elsevier Inc., 2004). doi:10.1016/B978-0-12-440905-7.50277-2

62. Grossman, K. J. & Stein, D. G. Does endogenous progesterone promote recovery of chronic sensorimotor deficits following contusion to the forelimb representation of the sensorimotor cortex? Behav. Brain Res. 116, 141–148 (2000).

63. Rubenstein, R. et al. Tau phosphorylation induced by severe closed head traumatic brain injury is linked to the cellular prion protein. Acta Neuropathol Commun 5, 30–17 (2017).

64. Farace, E. & Alves, W. M. Do women fare worse: a metaanalysis of gender differences in traumatic brain injury outcome. Journal of neurosurgery 93, 539–545 (2000).

65. Farin, A., Deutsch, R., Biegon, A. & Marshall, L. F. Sex-related differences in patients with severe head injury: greater susceptibility to brain swelling in female patients 50 years of age and younger. Journal of neurosurgery 98, 32–36 (2003).

66. Covassin, T., Elbin, R. J., Harris, W., Parker, T. & Kontos, A. The Role of Age and Sex in Symptoms, Neurocognitive Performance, and Postural Stability in Athletes After Concussion:. The American Journal of Sports Medicine 40, 1303–1312 (2012).

67. Tham, S. W. et al. Persistent Pain in Adolescents Following Traumatic Brain Injury. The Journal of Pain 14, 1242–1249 (2013).

68. Zuckerman, S. L. et al. Effect of sex on symptoms and return to baseline in sport-related concussion. J Neurosurg Pediatr 13, 72–81 (2014).

69. Wagner, A. K. et al. Evaluation of estrous cycle stage and gender on behavioral outcome after experimental traumatic brain injury. Brain Research 998, 113–121 (2004).

70. Maghool, F., Khaksari, M. & khachki, A. S. Differences in brain edema and intracranial pressure following traumatic brain injury across the estrous cycle: Involvement of female sex steroid hormones. Brain Research 1497, 61–72 (2013).

71. Fortress, A. M., Avcu, P., Wagner, A. K., Dixon, C. E. & Pang, K. C. H. Experimental traumatic brain injury results in estrous cycle disruption, neurobehavioral deficits, and impaired GSK3 β/β-catenin signaling in female rats. Experimental Neurology 315, 42–51 (2019).

72. Wagner, A. K., McCullough, E. H., of, C. N.J. 2011. Acute serum hormone levels: characterization and prognosis after severe traumatic brain injury. liebertpub.com 28, 871–888 (2011).

73. Hohl, A. et al. Luteinizing Hormone and Testosterone Levels during Acute Phase of Severe Traumatic Brain Injury: Prognostic Implications for Adult Male Patients. Front. Endocrinol. 9, 85–7 (2018).

74. Nance, E. A. et al. A Dense Poly(Ethylene Glycol) Coating Improves Penetration of Large Polymeric Nanoparticles Within Brain Tissue. Sci Transl Med 4, 149ra119–149ra119 (2012).

75. Smith, D. H. et al. A Model of Parasagittal Controlled Cortical Impact in the Mouse - Cognitive and Histopathologic Effects. J Neurotrauma 12, 169–178 (1995).

76. Yardeni, T., Eckhaus, M., Morris, H. D., Huizing, M. & Hoogstraten-Miller, S. Retro-orbital injections in mice. Lab Anim 40, 155–160 (2011).

77. Mostany, R. & Portera-Cailliau, C. A craniotomy surgery procedure for chronic brain imaging. JoVE – (2008). doi:10.3791/680

78. Holtmaat, A. et al. Long-term, high-resolution imaging in the mouse neocortex through a chronic cranial window. Nat Protoc 4, 1128–1144 (2009).

