## Supplementary files for "Sex-dependent macromolecule and nanoparticle delivery in experimental brain injury"

### Supplementary figures

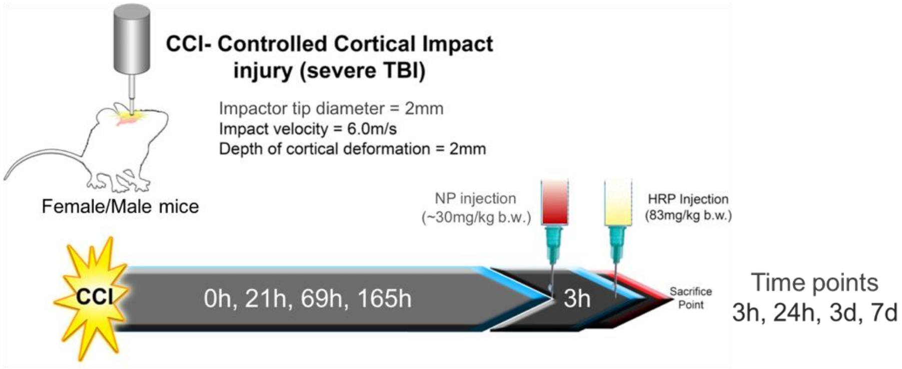


**Fig S1**: *In vivo* experimental study design: Nanoparticles (40 nm) were intravenously injected in CCI induced adult mice 3 h before sacrifice at 3 h, 24 h, 3 d and 7 d post-injury. The positive control BBB permeability marker, horseradish peroxidase (HRP) was injected 10 mins before sacrifice.

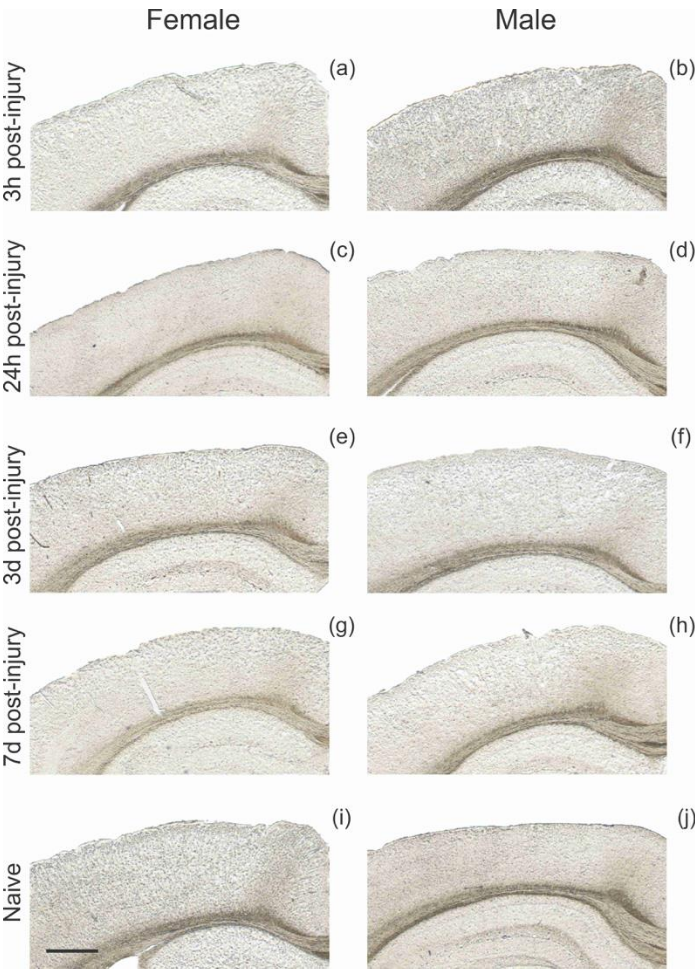


**Fig. S2**: HRP stained tissue after TBI in contralateral hemisphere: Representative images of HRP stained tissue at 3 h (a, b), 24 h (c, d), 3 d (e, f), 7 d (g, h) post-CCI and naïve (i, j). The first column shows the HRP response in female cohort and the second column in males. Scale bar=750μm.

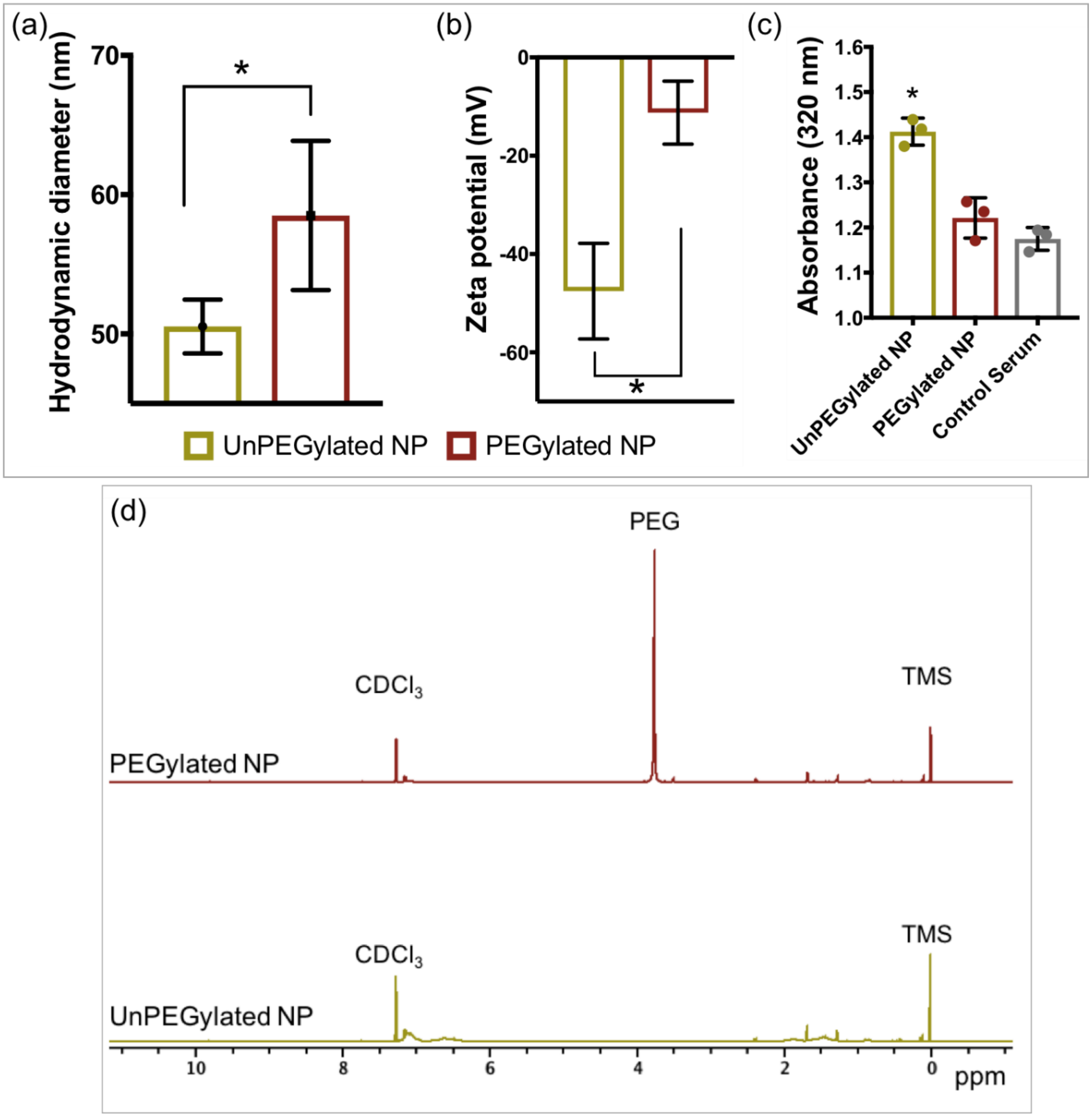


**Fig. S3: Nanoparticle characterization** (a) Hydrodynamic diameter of unPEGylated and PEGylated NPs, Student's t-test, * $p<0.05$, Error bars represent standard deviation, n=3. (b) Zeta potential of unPEGylated and PEGylated NPs. Student's t-test, * $p<0.05$, Error bars represent standard deviation, n=3. (c) Stability of NP: Absorbance of unPEGylated and PEGylated NP after 3 h incubation. UnPEGylated NPs shows significant increase in absorbance compared to control serum, * $p<0.05$, One-way ANOVA, Tukey's comparison. Error bars represent standard error of mean, n=3.

(d) $^1$H NMR of polystyrene NPs before and after PEGylation (2kDa). The appearance of PEG ($\partial$ = 3.63) peak confirms that the PEG was covalently linked to polystyrene NPs.

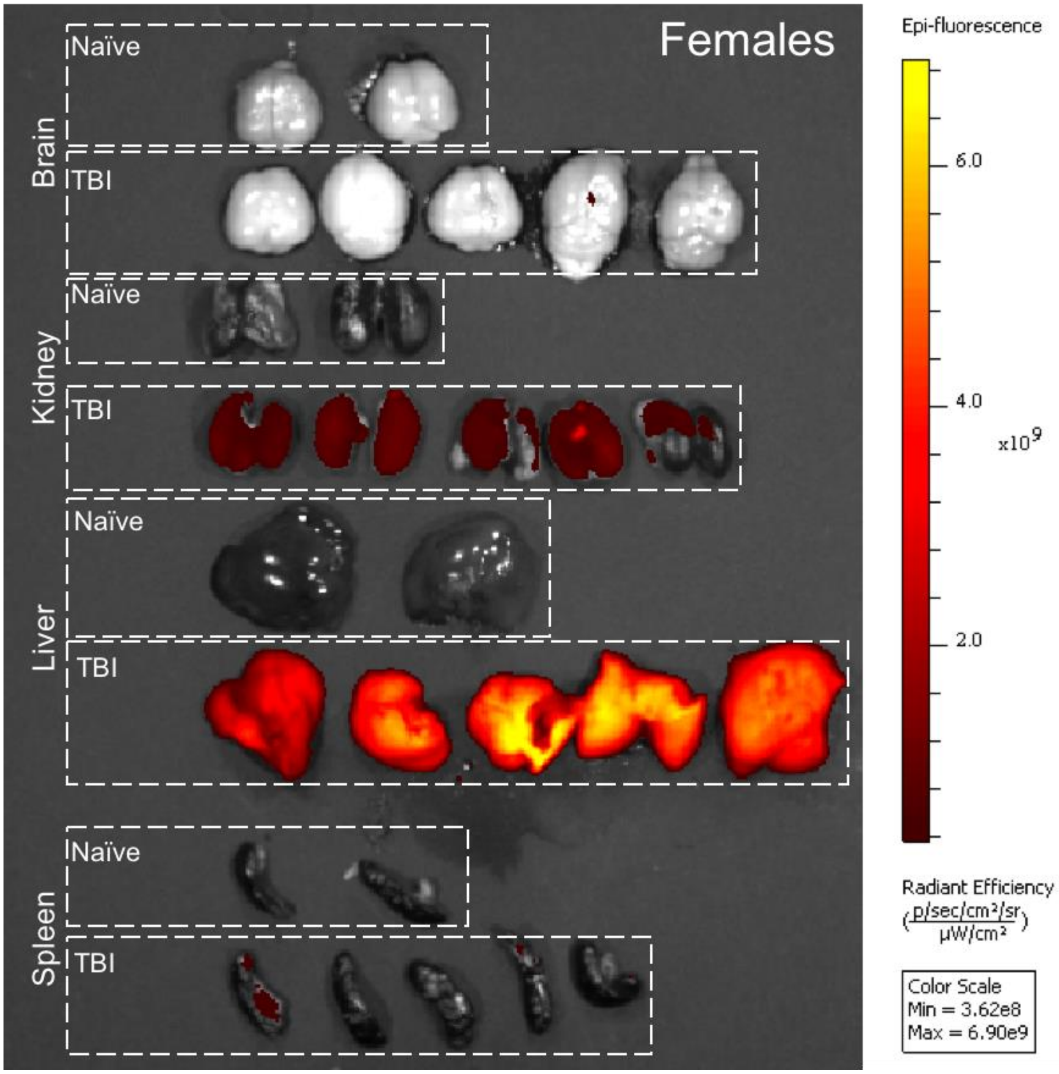


**Fig. S4: *Ex vivo* biodistribution of nanoparticles after 3 h post-injection.** (a) Ex vivo imaging of brain, kidney, liver and spleen of naïve animals and 7 d post-TBI animals. NP was intravenously injected 3 h before sacrifice. At the time of sacrifice (3 h after NP injection), terminal blood was drawn and plasma containing the NPs was isolated. Plasma concentration showed 30 % of the injected plasma concentration at 3 h post injection.

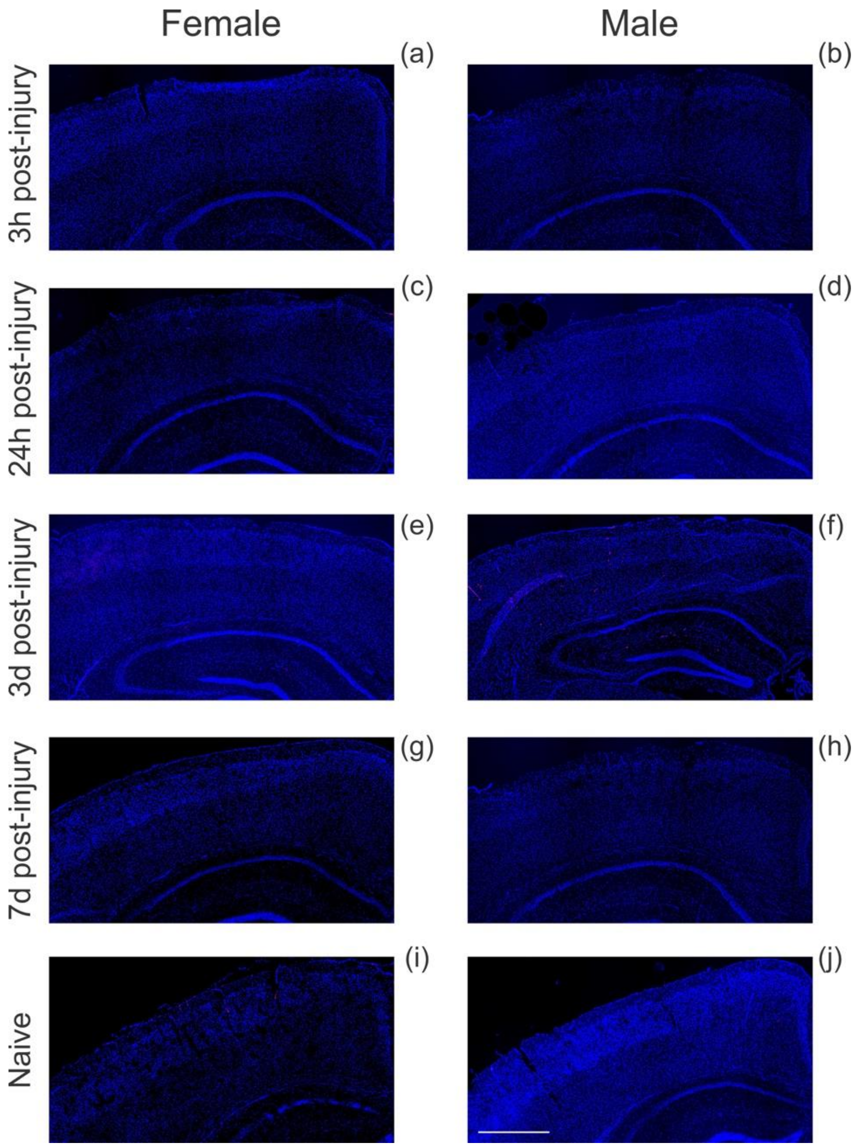


**Fig. S5: NP accumulation analysis after TBI in contralateral hemisphere:** Representative images of contralateral tissue at 3 h ((a)-(b)), 24 h ((c)-(d)), 3 d ((e)-(f)), 7 days ((g)-(h)) post-CCI and naïve ((i)-(j)). The first column shows female cohort and the second column in males. Scale bar = 750 μm.

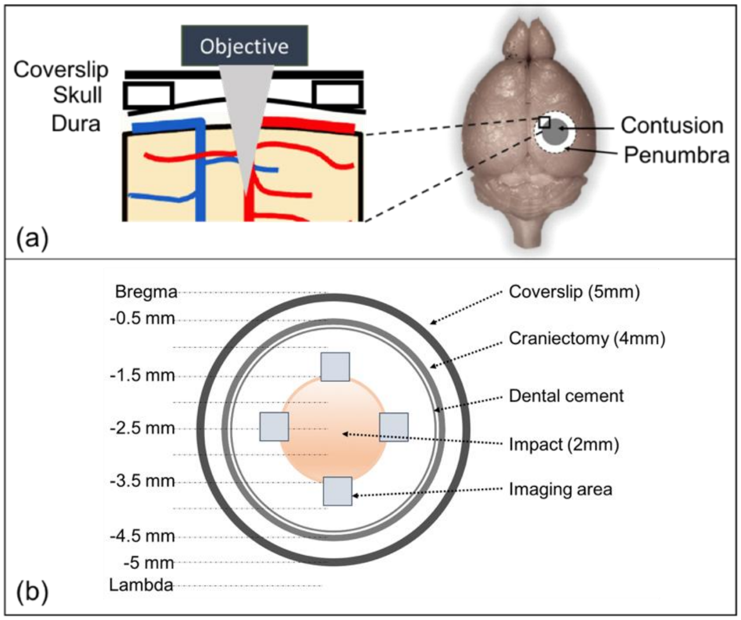


**Fig. S6: The experimental setup for two-photon microscopy imaging.** (a) Sagittal view of the entire setup. (b) Cross sectional view of the location of imaging region with respect to the contusion and penumbra region. Transgenic (CX3CR1-EGFP) female and male mice were used for this study. After a craniectomy (4 mm) and impact (2 mm), a glass coverslip of 5 mm was placed on top of the brain tissue and was secured with dental cement. A water-immersion objective lens was used and the anesthetized animal was secured on stereotaxic stage for two-photon imaging. Imaging area is shown as blue boxes located at the injury penumbra.

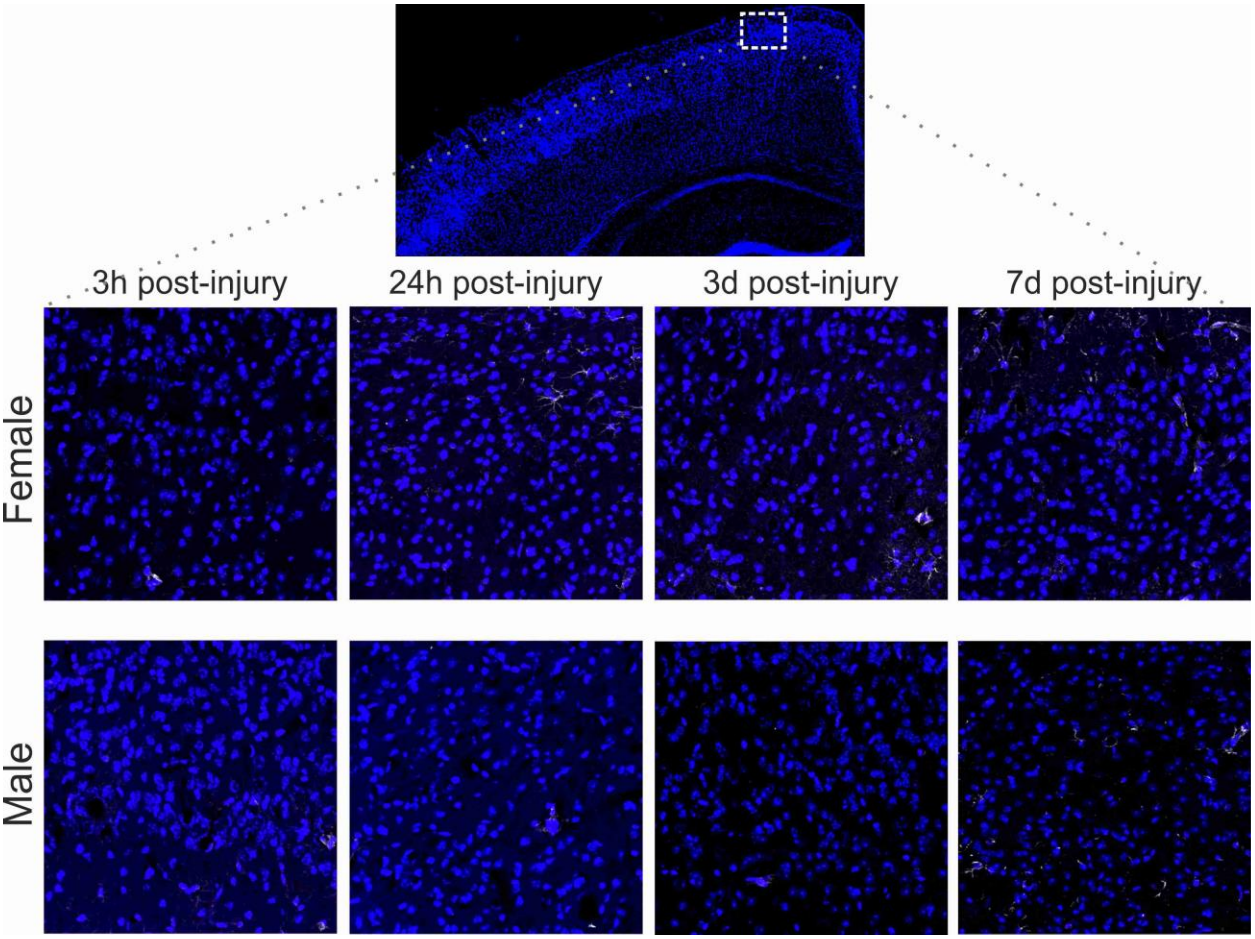


**Fig. S7:** Representative images of anti-glial fibrillary acidic protein (GFAP) staining in the contralateral hemisphere. Astrocyte reactivity at 3 h, 24 h, 3 d, and 7 d post-injury in females (top row) and males (bottom row) are shown in the image.

### Supplementary Tables

| | | Comparison | F (DFn, DFd) | P value | Significant |
|---|---|---|---|---|---|
| 1 | HRP: Female cohort | Interaction | F (3, 24) = 34.35 | P<0.0001 | Yes |
| | | Time points | F (3, 24) = 34.14 | P<0.0001 | Yes |
| | | Ipsi vs Contra | F (1, 24) = 297.4 | P<0.0001 | Yes |
| 2 | HRP: Male cohort | Interaction | F (3, 24) = 33.46 | P<0.0001 | Yes |
| | | Time points | F (3, 24) = 34.4 | P<0.0001 | Yes |
| | | Ipsi vs Contra | F (1, 24) = 177.7 | P<0.0001 | Yes |
| 3 | HRP: Female vs Male cohort (ipsilateral) | Interaction | F (3, 24) = 4.519 | P=0.0119 | Yes |
| | | Time points | F (3, 24) = 67.64 | P<0.0001 | Yes |
| | | Sex | F (1, 24) = 17.23 | P=0.0004 | Yes |

**Supplementary Table 1.** Statistical analysis results of HRP extravasation: Tabular results of two-way ANOVA of extravasation of intravenously injected macromolecular tracer (horseradish peroxidase) after focal brain injury. Row 1 shows the two-way ANOVA results of female cohort, row 2 shows the results of male cohort and row 3 shows the result of ipsilateral hemisphere comparison of female and male cohort.

| | | Comparison | F (DFn, DFd) | P value | Significant |
|---|---|---|---|---|---|
| 1 | NP: Female cohort | Interaction | F (3, 24) = 23.19 | P<0.0001 | Yes |
| | | Time points | F (3, 24) = 22.73 | P<0.0001 | Yes |
| | | Ipsi vs Contra | F (1, 24) = 100.6 | P<0.0001 | Yes |
| 2 | NP: Male cohort | Interaction | F (3, 24) = 128.5 | P<0.0001 | Yes |
| | | Time points | F (3, 24) = 134.1 | P<0.0001 | Yes |
| | | Ipsi vs Contra | F (1, 24) = 236.5 | P<0.0001 | Yes |
| 3 | NP: Female vs Male cohort (ipsilateral) | Interaction | F (3, 24) = 4.876 | P=0.0087 | Yes |
| | | Time points | F (3, 24) = 94.85 | P<0.0001 | Yes |
| | | Sex | F (1, 24) = 1.45 | P=0.2402 | No |

**Supplementary Table 2.** Statistical analysis results of NP accumulation: Tabular results of two-way ANOVA of intravenously injected NP accumulation after focal brain injury. Row 1 shows the two-way ANOVA results of female cohort, row 2 shows the results of male cohort and row 3 shows the result of ipsilateral hemisphere comparison of female and male cohort.

| Comparison | F (DFn, DFd) | P value | Significant |
|---|---|---|---|
| Interaction | F (3, 32) = 0.3234 | P=0.8084 | No |
| Time points | F (3, 32) = 9.445 | P=0.0001 | Yes |
| Sex | F (1, 32) = 11.68 | P=0.0017 | Yes |

**Supplementary Table 3.** Statistical analysis results of T2 weighted MRI lesion volume: Tabular results of two-way ANOVA of lesion volume (# voxels) in the ipsilateral cortex comparisons between sexes after focal brain injury.

| | | Comparison | F (DFn, DFd) | P value | Significant |
|---|---|---|---|---|---|
| 1 | GFAP: Female cohort | Interaction | F (3, 24) = 6.332 | P=0.0026 | Yes |
| | | Time points | F (3, 24) = 5.605 | P=0.0047 | Yes |
| | | Ipsi vs Contra | F (1, 24) = 19.99 | P=0.0002 | Yes |
| 2 | GFAP: Male cohort | Interaction | F (3, 24) = 10.15 | P=0.0002 | Yes |
| | | Time points | F (3, 24) = 11.94 | P<0.0001 | Yes |
| | | Ipsi vs Contra | F (1, 24) = 28.96 | P<0.0001 | Yes |
| 3 | GFAP: Female vs Male cohort (ipsilateral) | Interaction | F (3, 24) = 0.0868 | P=0.9666 | No |
| | | Time points | F (3, 24) = 15.63 | P<0.0001 | Yes |
| | | Sex | F (1, 24) = 0.2255 | P=0.6391 | No |

**Supplementary Table 4.** Statistical analysis results of percentage area of GFAP staining after focal brain injury. Tabular results of two-way ANOVA of percent area GFAP positive staining comparisons between sexes after focal brain injury. Row 1 shows the two-way ANOVA results of female cohort, row 2 shows the results of male cohort and row 3 shows the result of ipsilateral hemisphere comparison of female and male cohort.

### Supplementary methods

#### *Ex vivo* biodistribution and blood plasma concentration

A separate cohort was used for biodistribution and plasma concentration analysis. At 7 d post-injury animals were intravenously injected with 50 µl NPs and were sacrificed after 3 h post-injection. As a control group un-injected animals were sacrificed. Whole blood, brain, kidney, liver and spleen were harvested for further analysis from TBI and control animals. Fluorescent images were acquired under the IVIS Imaging System (Xenogen Imaging Technologies, Alameda, CA, USA) at Ex $\lambda = 570\ nm,$ Em $\lambda = 620\ nm.$. Plasma was collected from the blood and the total NP concentration was analyzed using a standard curve obtained from serial dilution of the PEGylated NPs. The plasma samples were read using a fluorescent plate reader (BioTek Instruments, Inc., Winooski, VT, USA).
