## Supplementary figures and images for "Sex-dependent macromolecule and nanoparticle delivery in experimental brain injury"

### Sham Cohort

#### Sham Cohort

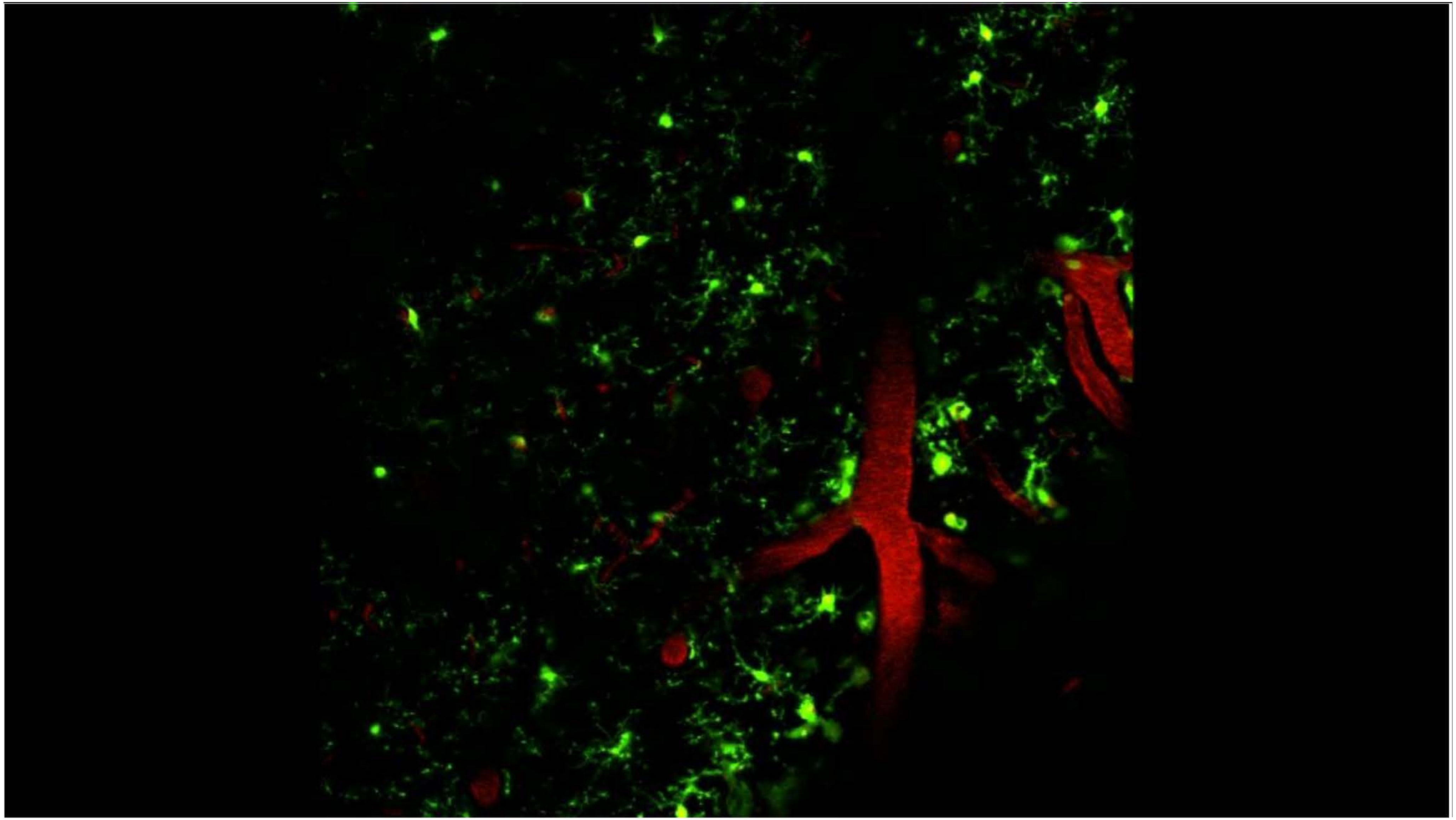

### Two-Photon Videos Female Cohort

#### Female Cohort – All time points

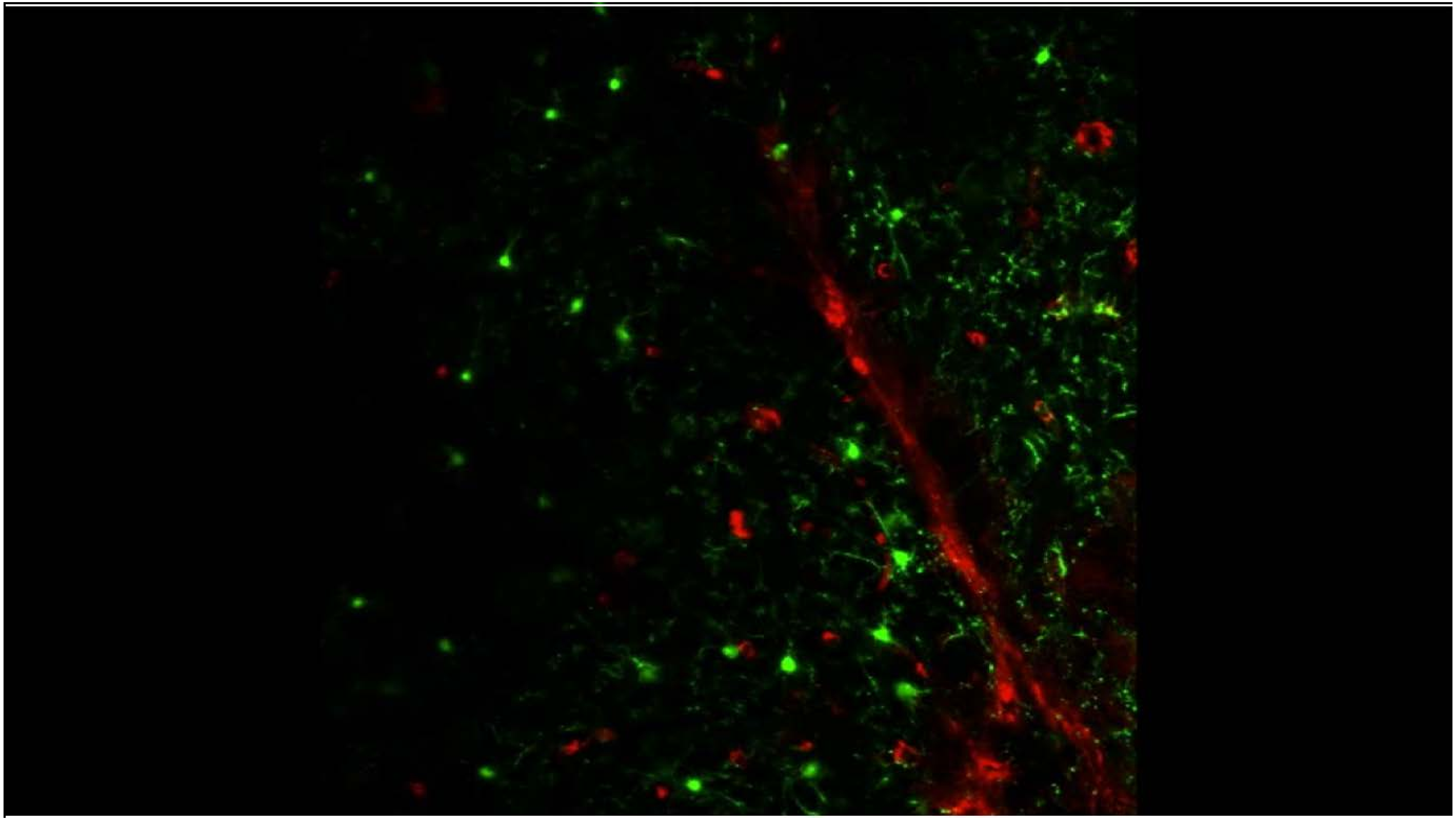

### Two-Photon Videos Male Cohort

#### Male Cohort – All time points

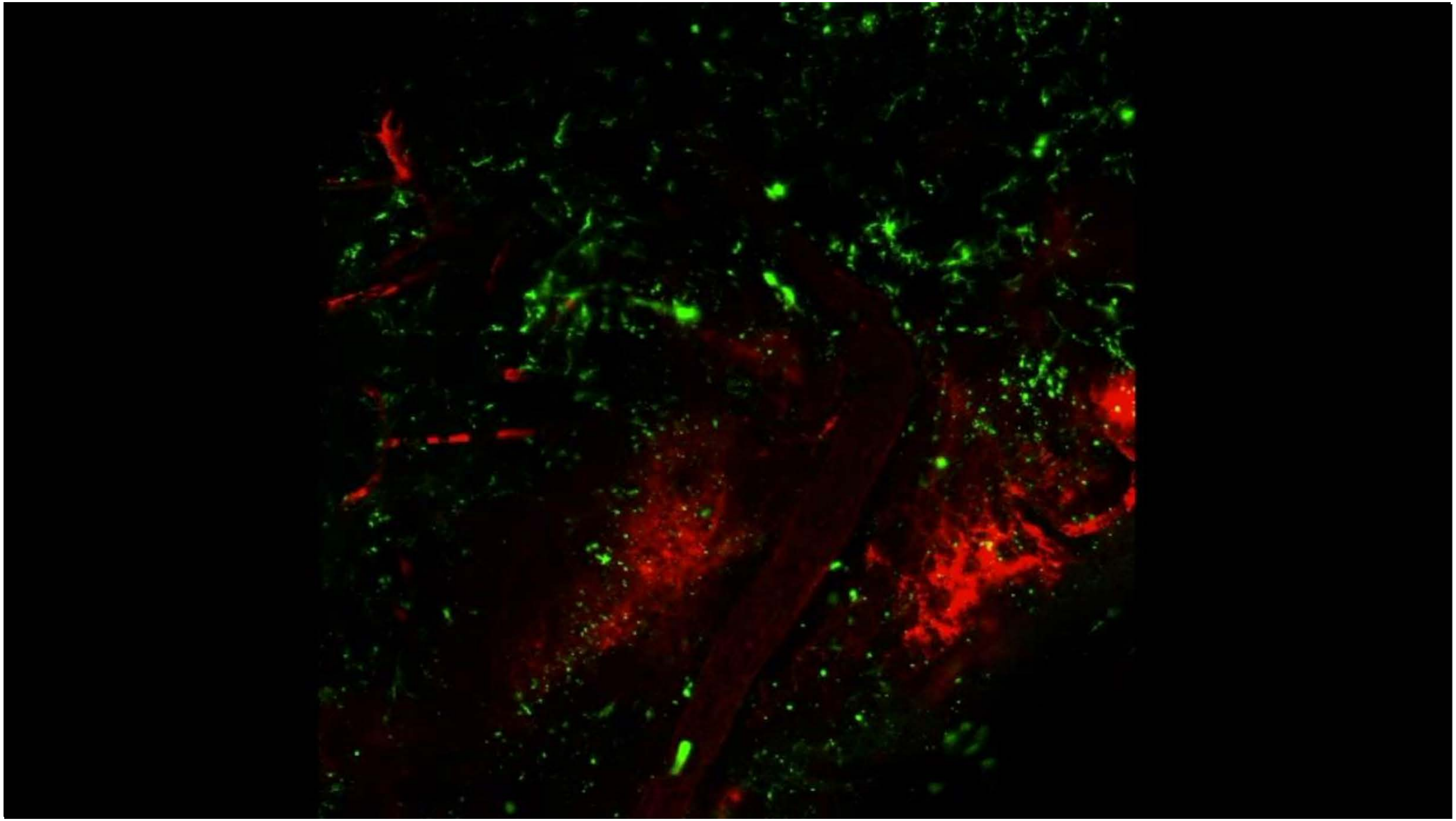
